# Hyperactive 20S Proteasome Enhances Proteostasis and ERAD in *C. elegans* via degradation of Intrinsically Disordered Proteins

**DOI:** 10.1101/2024.04.04.588128

**Authors:** David Salcedo-Tacuma, Nadeem Asad, Raymond Anderson, David M. Smith

## Abstract

Age-related proteinopathies, including Alzheimer’s and Parkinson’s disease, are driven by the toxic accumulation of misfolded proteins, particularly intrinsically disordered proteins (IDPs), that overwhelm cellular proteostasis. The proteasome is responsible for the clearance of these proteins, but it is unclear why it fails to do so in these diseases. Here, we report a novel strategy employing a *C. elegans* model with a hyperactive 20S proteasome (α3ΔN) to achieve selective activation. This activation robustly enhances the degradation of IDPs and misfolded proteins, markedly reduces oxidative damage, and significantly improves ER-associated degradation (ERAD). Notably, aggregation-prone substrates, such as endogenous vitellogenins and human alpha-1 antitrypsin (ATZ), are efficiently cleared. Proteomic and transcriptomic reprogramming reveals systemic adaptations characterized by increased protein turnover and enhanced oxidative stress resistance, independent of superoxide dismutases. Strikingly, proteasome hyperactivation extends lifespan and enhances stress resistance independently of known proteostasis pathways including the canonical unfolded protein response mediated by xbp-1. Our findings provide substantial support for a “20S pathway” of proteostasis that alleviates protein aggregation and oxidative stress, offering a promising therapeutic strategy for neurodegenerative diseases

**Teaser:** Enhanced disordered protein clearance by the 20S proteasome in worms prevents toxic protein buildup and promotes stress resistance and longevity.

## Introduction

Age-related proteinopathies, such as Alzheimer’s and Parkinson’s disease, pose a significant challenge to public health due to their devastating impact on the aging population. These disorders are characterized by the accumulation of misfolded proteins (1–7), particularly intrinsically disordered proteins (IDPs) which lack stable three-dimensional structures and are prone to aggregation (1–7). While IDPs play key roles in cellular regulation and signaling, their structural flexibility makes them particularly vulnerable to misfolding, driving disease progression through the accumulation of toxic protein species that lead to neuronal dysfunction.

The cellular proteostasis network, a complex and interconnected system, maintains protein homeostasis through the coordination of protein synthesis, folding, and degradation (1, 8–10). Molecular chaperones facilitate the folding of newly synthesized proteins, refold misfolded proteins, and prevent protein aggregation (8, 10–12). These chaperones work in concert with degradation pathways, such as autophagy, ER Associated Degradation (ERAD), and the ubiquitin-proteasome system (UPS), to remove damaged or misfolded proteins and prevent the accumulation of proteotoxic species (8, 10–12). The proteostasis network also integrates stress response pathways, including the oxidative stress response, heat shock response (HSR), and unfolded protein response (UPR), which comprise stress indicators allowing the network to adapt to changing cellular conditions (13, 14). Disruptions to this finely balanced system can have far-reaching consequences, contributing to the development and progression of various diseases, particularly those associated with aging (10, 11, 15).

The ubiquitin-proteasome system (UPS) is central to proteostasis, targeting ubiquitinated proteins for breakdown by the 26S proteasome, which consists of a 20S core particle and 19S regulatory subunits. However, the 20S proteasome can function independently of ubiquitin and the 19S regulator, offering a specialized pathway for degrading IDPs and oxidatively damaged proteins. Notably, certain pathological oligomers, such as those derived from amyloid-β, α-synuclein, and Huntingtin, have been shown to inhibit proteasome activity by allosterically inhibiting the opening of the 20S proteasome gate (16, 17). The open 20S proteasome is highly capable of degrading IDPs since unfolded proteins can diffuse into the opened 20S. In contrast, folded proteins typically require ubiquitination and targeting to the 26S proteasome (19S-20S), which can unfold these substrates to inject them into the 20S core (18, 19).

Consequently, the ability of the 20S proteasome to independently degrade misfolded proteins is particularly relevant in aging and neurodegenerative conditions, where proteasome pathways are often disrupted. Enhancing proteasome activity emerges as a promising strategy for combating age-related proteinopathies. For example, specific HbYX motif peptides have been shown to facilitate and enhance the 20S proteasome’s ability to degrade IDPs and reverse the inhibitory effects of toxic oligomers in-vitro (17, 20, 21). Additionally, pharmacological upregulation of the 26S proteasome has been shown to improve the clearance of misfolded proteins (22). Similarly, genetic overexpression of 20S proteasome subunits in several animal models has proven beneficial, leading to enhanced aggregation-prone/misfolded protein degradation, extended lifespan, and increased resistance to stressors (23–29). Although overexpression-based models have demonstrated benefits, they fail to address whether intrinsic changes to 20S proteasome activity can unlock distinct degradation mechanisms and substrate preferences, potentially uncovering novel aspects of proteasome function and substrate targeting.

Given the ubiquitous presence of the UPS across diverse tissues and its involvement in numerous cellular processes, elucidating the specific roles of the independent 20S proteasome is inherently challenging, since current proteasome inhibitors target the 20S and thus affect all proteasome forms—including the 26S. Thus, to determine the therapeutic potential of the 20S proteasome, we selectively hyper-activated it. This targeted approach represents the most effective, perhaps only, strategy to explore 20S function in biology.

In this study, we use the nematode Caenorhabditis elegans, a powerful model organism due to its short lifespan, genetic flexibility, and conserved proteostasis pathways. A hyperactive 20S proteasome model was developed by using CRISPR-Cas9 to induce a truncation of the N-terminal domain of the α3 subunit (pas-3) of the 20S proteasome (α3ΔN-20S). This N-terminal truncation selectively generates a constitutively open-gate form of the 20S (28). Throughout this manuscript, we refer to this open gate mutant strain pas-3(dsm100) as α3ΔN. This mutation is considered hyperactivating since it induces a continuously activated form of the 20S that is physiologically distinct from the normal activated state of the 20S. This α3ΔN *C. elegans* strain differs from all other prior “proteasome enhancement” animal models available to date, which are based on proteasomal subunit overexpression methods (e.g., increased proteasome amount) or pharmacological approaches that alter signaling pathways effecting 26S regulation.

While the basic phenotypic characterization of the α3ΔN mutant demonstrated notable benefits, such as extended lifespan, heat shock resistance, and a specific 250-fold activation of the 20S proteasome activity relative to 26S activity (28), the underlying mechanisms linking 20S activation to these phenotypic effects and their potential therapeutic implications were not addressed. Additionally, this constitutively active proteasome is expected to affect the typical degradation rates of some proteins, especially nascent proteins and IDPs, although this has not been determined. As most pathological proteins that accumulate in neurodegenerative diseases are IDPs (30–35), the disorder-dependent “20S pathway” for degradation becomes highly relevant in the context of the α3ΔN mutant’s hyperactivation.

Using combined proteomic and transcriptomic analyses, we examine how this hyperactivation of the 20S proteasome reshapes the proteome, improves protein quality control, and increases resistance to both oxidative stress and endoplasmic reticulum (ER) stress. The ER-associated degradation (ERAD) pathway, which clears misfolded proteins from the ER, is vital for proteostasis, and its failure is implicated in diseases like alpha-1 antitrypsin deficiency. We demonstrate that proteasome hyperactivation in *C. elegans* leads to the selective degradation of IDPs, which is associated with increased protein synthesis. Our hyperactive strain showed decreased oxidative damage and was even resistant to oxidative stresses in the absence of the SOD system the cell’s primary defense against oxidative damage. Additionally, while the 26S is a known endpoint mediator of ERAD, we found that 20S proteasome hyperactivation substantially enhanced ERAD, as evidenced by the efficient clearance of aggregation-prone ER resident proteins such as VIT-2 and ATZ. VIT-2 is an ApoB homolog and known endogenous ERAD substrate, while ATZ is also an ERAD substrate in humans that accumulates in the liver in alpha-1 antitrypsin deficiency (36, 37).

Our key mechanistic finding reveals that the 20S proteasome hyperactivation has a profound capacity to enhance oxidative stress resistance and ERAD function leading to increased lifespan independent of the UPR. Taken together, this study demonstrates that activation of IDP degradation in *C. elegans* by 20S hyperactivation is an independent mechanism for direct proteostasis management and longevity enhancement. The targeted degradation of IDPs that form aggregates highlights the potential of proteasome hyperactivation to alleviate the accumulation of toxic protein aggregates in the context of proteinopathies and age-related diseases, of which there are many. Our results reveal the 20S proteasome degradation pathways as a regulator of proteostasis and underscore its potential as a therapeutic target for diseases driven by protein misfolding and aggregation. By selectively boosting 20S activity, it may be possible to eliminate harmful proteins while preserving overall protein balance, offering a precise strategy for tackling proteinopathies and laying the groundwork for new ways to combat cellular stress and neurodegenerative diseases linked to aging.

## Results

### Proteome reconfiguration and adaptive protein turnover following 20S proteasome hyperactivation

We assessed the overall proteomic changes in the α3ΔN mutant using Tandem Mass Tag-mass spectrometry (TMT-MS). We focused on 8-day old adult worms (n=5), a post-reproductive stage that allow us to compare against wild type without the confounding effects from decreased egg numbers in the α3ΔN mutant compared to WT. In addition, FUDR was used in both populations to normalize any physiological effects due to differences in fecundity, as was done in (Anderson et. al.(28)). Furthermore, this stage marks the onset of age-related phenotypes and detectable age-related proteomic and transcriptomic signatures in *C.elegans* (11, 38, 39). We identified 4,267 proteins (Q-value<0.01), and unsupervised hierarchical clustering combined with Principal Component Analysis (PCA) revealed distinct proteomic profiles between the α3ΔN mutant and wild type (Fig. 1A). Using DEqMS (40) (FDR < 0.05, |Log2FC| > 0.59), we detected 459 differentially expressed proteins (DEPs), representing approximately 10% of the total proteome (Table S1). Among these, 252 were downregulated and 207 upregulated in the α3ΔN mutant (Fig. 1B-FigS1A).

**Figure 1.**
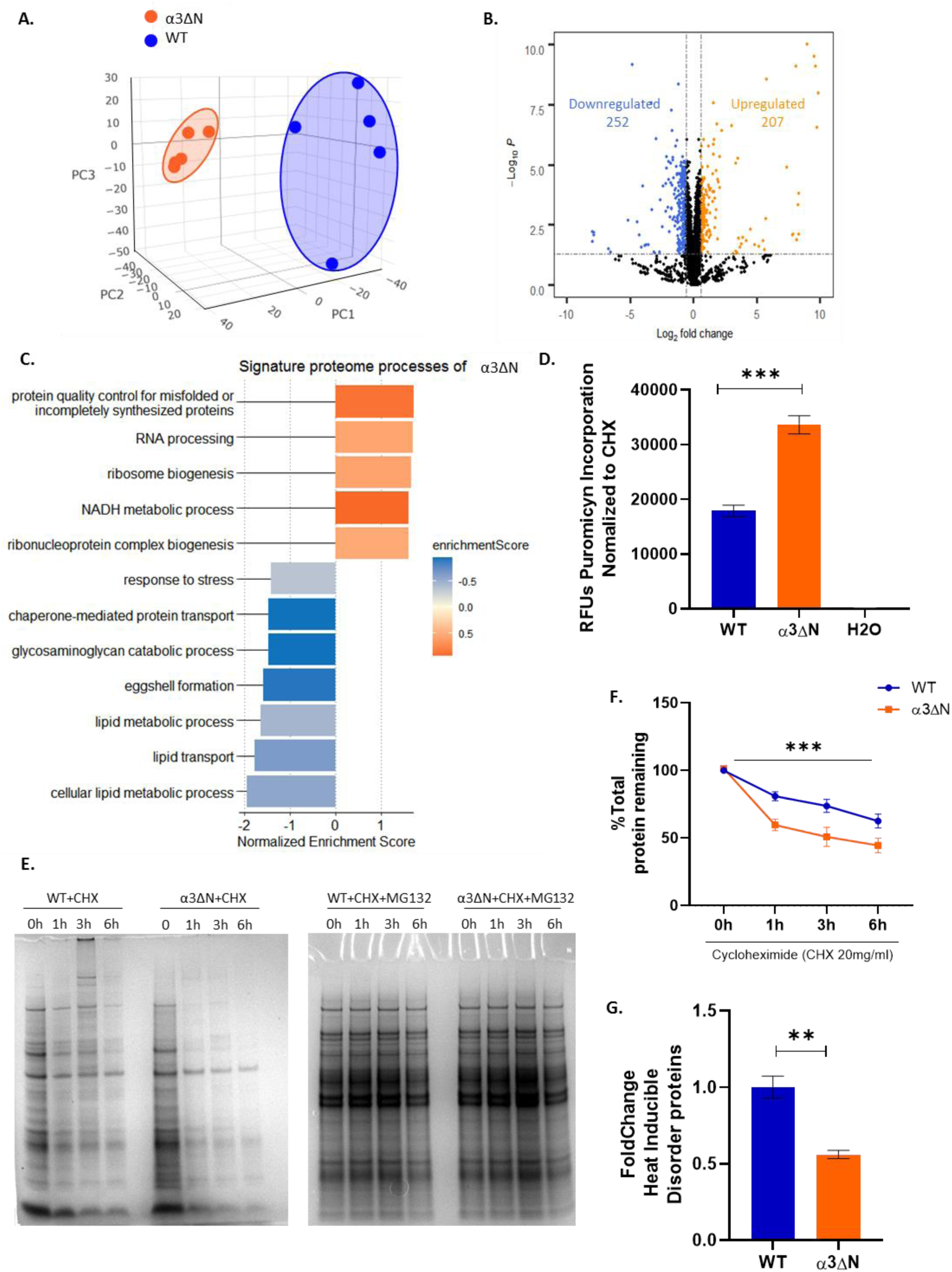
Open gate mutant α3ΔN rewires the proteome of *C. elegans decreasing disorder* proteins and increasing global synthesis: **A)** Dimensionality reduction via three-dimensional PCA was performed on TMT-MS data to depict profile differences between n=5 wild-type (WT) and α3ΔN mutants. **B,** Volcano plot of Differentially Expressed Proteins (DEPs) detected Log2FC α3ΔN/WT vs - log10FDR. All points above the dotted line are statistically significant (FDR<0.05). Blue points above the x axis dotted line represent DEPs downregulated (FC<-1.5) and orange dots represent DEPs upregulated (FC>1.5). **C.** GSEA of DEPs found in α3ΔN mutants, x axis represents the Normalized Enriched Score for each of the categories according to ClusterProfiler in R; FDR<0.05. **D.** Rates of protein translation in live *C.elegans* determined by 6-FAM-puromycin, lysing and, separating free 6-FAM-puro from labeled nascent proteins with desalting columns, following by fluorescent quantification in plate reader. Bars represent mean +/-SEM of n=6, \*\*\**p<0.01 ordinary One-way-ANOVA*. E-F. *CHX* Pulse-chase assay to assess global protein degradation rates over 6 h of CHX or CHX+MG132 treatments in α3ΔN mutants. CHX/MG132 ratios were normalized to the 0h timepoint of the respective strain. **(F)** Quantification of E. Values are depicted as mean ± SEM generated from *n* = 3 independent experiments. Statistical significance was determined using Satterthwaite’s degrees of freedom method with ImerTest, ***p<0.001 indicating a statistically significant difference between conditions. **G.** Quantification of intrinsically Disorder Proteins (IDPs) content in WT and α3ΔN mutants. IDPs from lysates were enriched by two steps of heat denaturation and precipitation of folded proteins, leaving primarily IDPs. SDS-page gels were quantified after commassie staining. Bars represent means +/-SEM. n=3, \*\**p<0.01 unpaired t-test*

Next, to elucidate the biological significance of the proteome profile changes, we performed Gene Set Enrichment Analysis (GSEA) (41). The analysis identified two primary functional groups affected by proteasome hyperactivation: protein quality control, and lipid metabolism processes (Fig1C-FigS1B). Among the upregulated processes we detected a notable enrichment related to folding and translational activity, including ribosome biogenesis, RNA-Binding Proteins (RBPs), and RNA processing. In normal physiological conditions, the proteasome mediates the degradation of DNA and RNA binding proteins, which are enriched in intrinsic disorder regions (IDRs). This is essential for maintaining appropriate protein levels for vital cellular functions such as gene expression, DNA repair, and RNA processing (42–44).

The upregulation of RBPs, DNA and RNA binding proteins in the α3ΔN mutants suggests a selective degradation pressure on these substrates, which the organism may compensate for by upregulating these critical proteins to high levels in order to preserve these vital functions. This scenario aligns with the hypothesis that enhanced degradation of IDPs, facilitated by the α3ΔN-20S, leads to a rapid turnover of certain proteins and potentially drives a higher rate of protein synthesis. To test this hypothesis, we labelled newly synthesized proteins in live worms using 6-FAM-puromycin (45). To ensure the accuracy of our measurements, we normalized the fluorescence signals to those obtained from worms treated with cycloheximide, a well-known inhibitor of protein biosynthesis. We observed a nearly 2-fold increase in protein synthesis in the α3ΔN mutant (Fig1D), validating our hypothesis. This acceleration in synthesis is likely crucial for sustaining proteome stability in the face of accelerated degradation by the α3ΔN proteasome illustrating a sophisticated mechanism by which cells adapt to ensure continued function during the increased protein turnover.

To test if the α3ΔN mutant had enhanced protein degradation in the α3ΔN strain, we performed a cycloheximide pulse chase assay. We blocked new protein synthesis with cycloheximide and measured the decrease of total protein levels by SDS-gel in the presence or absence of MG132 a well-known proteasome inhibitor which allowed us to normalize the degradation from proteasome solely. We observed that the α3ΔN mutant degraded proteins significantly more rapidly than WT (Fig 1E-F). The simultaneous acceleration of protein degradation and protein synthesis supports the hypothesis that enhanced protein synthesis is likely an adaptive response to balance out proteasome hyperactivation to maintain proteome stability.

We next investigated the impact of the hyperactive proteasome on degradation of intrinsically disordered proteins (IDPs), which can be degraded by the 20S in a ubiquitin-independent manner (30, 42, 46). To specifically assess the impact of proteasome hyperactivation on IDPs, we employed a controlled heat treatment protocol, as described in prior studies (3), to selectively enrich for heat-stable proteins. This approach effectively isolates a subset of proteins predominantly comprising IDPs, allowing us to focus on their degradation dynamics. Using this method, our analysis (Fig. 1G) revealed a striking 50% reduction in heat-stable protein levels in the α3ΔN mutant compared to WT, demonstrating significantly accelerated degradation of this protein pool. These results highlight how proteasome hyperactivation directly contributes to proteostasis by efficiently eliminating IDPs, which are often associated with age related protein and cellular dysfunction.

### Coordinated transcriptomic-proteomic signatures highlight broad proteostasis reprogramming in α3ΔN mutants

Given the significant alterations in protein synthesis, degradation, and translation observed in α3ΔN mutants, we hypothesized significant transcriptomic rewiring as part of a response to proteasome hyperactivation. To explore this, we conducted RNA sequencing on 8-day-old adult worms (n=5). PCA analysis (Fig. 2A) revealed distinct transcriptomic profiles for the α3ΔN mutant and wild type strains, mirroring the proteomic differences and suggesting a broad reprogramming of molecular pathways due to proteasome hyperactivity. Differential expression analysis identified 1,756 significantly altered genes in α3ΔN mutants (TableS2-FDR<0.05), with 854 upregulated and 902 downregulated (Fig.2B, FigS2A). Importantly, 20S proteasomal subunit expression remained unchanged (Fig.S2B-C), confirming that enhanced proteasomal activity is attributed solely to the open-gate modification rather than altered proteasome abundance.

**Figure 2.**
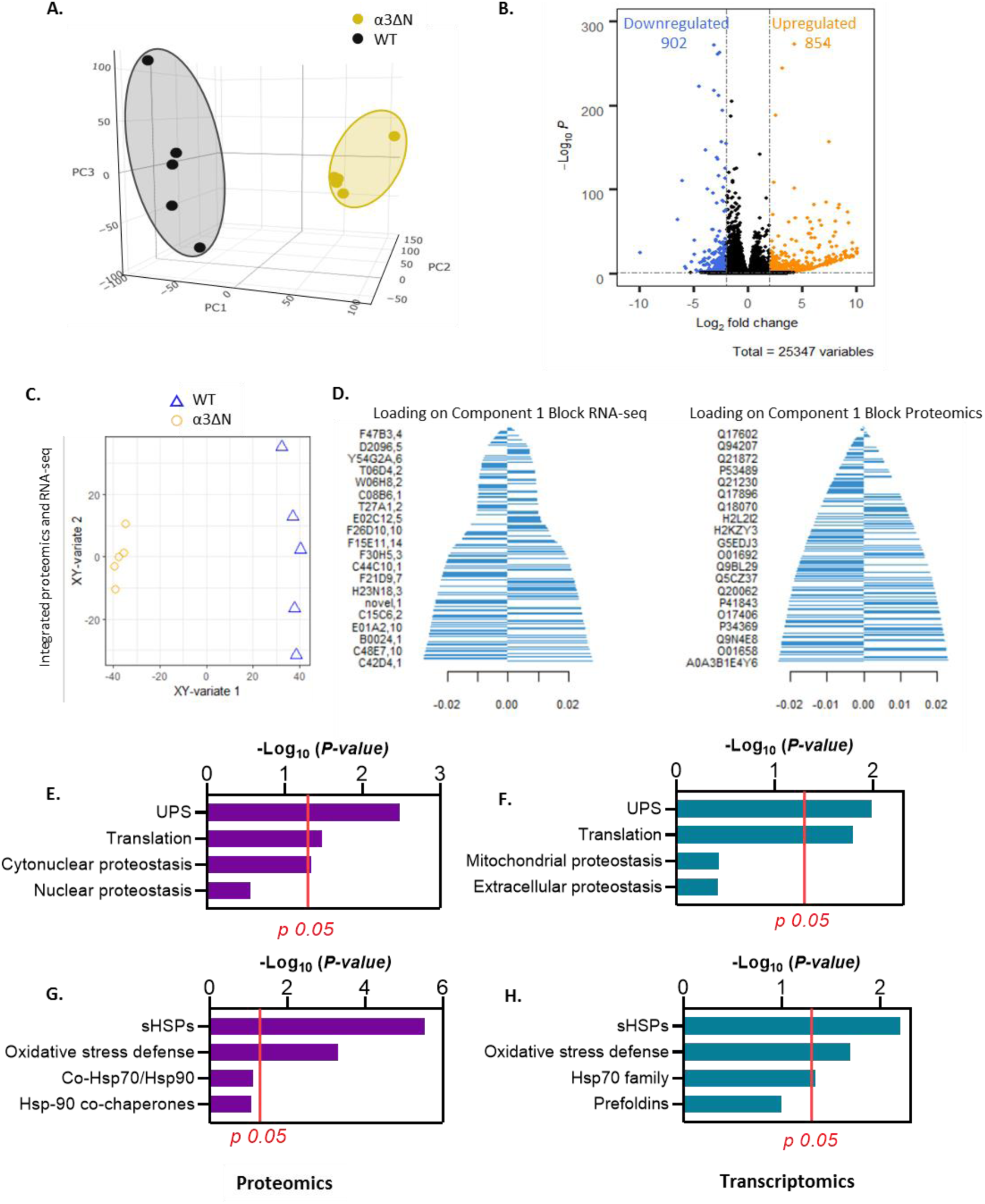
Open gate mutant α3ΔN impacts the transcriptome of proteostasis components involved in oxidative stress defense. **A.** Three Dimensional PCA of RNA-seq data to depict differences in the expression profile of WT and α3ΔN mutants (n=5). **B,** Volcano plot of Differentially Expressed Genes (DEGs) detected Log2FC α3ΔN/WT vs - log10FDR. All points above the dotted line are statistically significant (FDR<0.05). Blue points above the x axis dotted line represent DEGs downregulated (FC<-2) and orange dots represent DEGs upregulated (FC>2). **C.** Integration of proteomics and RNAseq data from α3ΔN samples using sparse projection to latent structures (sPLS). Samples are color-coded and shape-coded according to specific conditions, illustrating their projection within a combined integrated space. This visualization highlights distinct clusters, demonstrating similarities and differences across conditions. **D.** Barplots of Loadings. The left barplot shows the loadings of RNA transcripts, and the right barplot displays the loadings of proteins in the first sparse projection to latent space variable that differentiates between the α3ΔN and WT conditions. These barplots highlight the key proteins and transcripts and their relative importance in distinguishing between these conditions **E-F.** GSEA analysis of Proteostasis Network components was performed on proteomic (**E**) and transcriptomic (**F**) datasets using ClusterProfiler in R. Significantly enriched components (*adjusted p-value < 0.05*) are displayed, with the x-axis representing *–log10(p-value)* and the red line marking the significance threshold (–log10(0.05) = 1.3). **G-H.** Enrichment Analysis of the Chaperone subnetwork using the Hypergeometric Distribution for both omic datasets with *adjusted p-value* of <0.05 indicating significant enrichment. The x-axis indicates the –log10(*p*-value) and red line indicates a threshold of *adj-p* = 0.05 [−log10(0.05) =1.3].

To elucidate how changes in gene expression and protein production are interconnected in α3ΔN mutants—revealing a coordinated molecular response to proteasome hyperactivity that would be overlooked by examining either dataset independently—we employed data integration. Using sparse Partial Least Squares (sPLS) in mixOmics (47) we combined proteomic and transcriptomic datasets, as detailed in the Methods. The integrated analysis (Fig. 2C) revealed two distinct clusters corresponding to the α3ΔN and wild-type genotypes. This clear separation corroborates our previous findings, confirming that these genotypes exhibit distinct molecular profiles. Notably, the structural patterns observed in the sPLS analysis are consistent with the principal component analysis (PCA) structural profiles of the individual proteomic and RNA-seq datasets (Figs. 1A-2A), further validating these results. Importantly, the data integration revealed that most genes and proteins contributed modestly to the observed separation, suggesting that the response in the α3ΔN strain is not driven by a small subset of genes or proteins (Fig. 2D). Instead, we observe a systemic shift across the entire genetic and proteomic landscape (Figs. S3A-B, Table S6). Furthermore, analysis of transcription factor binding profiles using JASPAR indicated the involvement of numerous transcription factors (TFs) (Fig. S3C-TableS3), supporting the notion of a network-wide alteration in the α3ΔN strain. Collectively, these results point to a broad, interconnected adaptation rather than isolated changes, highlighting the complex nature of the biological processes affected by the α3ΔN mutation (Figs. S1B-S2D). This integrative approach allowed us to focus on system-level responses, which are crucial for a mechanistic understanding of the biological effects associated with a hyperactive proteasome.

Understanding the systemic adaptations underlying proteasome hyperactivation requires identifying the specific components driving these changes. Given the shared enrichment patterns observed across both omics datasets, we focused on identifying whether this systemic adaptation involved key elements of the proteostasis network a critical regulator of cellular homeostasis. To do so, we utilized pre-existing annotated families of Proteostasis Network components (PN) (48) and the chaperone subnetwork families from *C. elegans* (49) and performed enrichment analyses on both omic datasets. GSEA analysis on PN components revealed significant enrichment of UPS related proteins and translation components across both the proteomic and trancriptomic datasets, with an additional enrichment of cytonuclear proteostasis observed specifically in the proteomic dataset (Fig. 2E-F). This enrichment captures the broad reprogramming of proteostasis in our study, wherein increased translation (Fig 1D) and enhanced UPS-mediated degradation (Fig 1E-G) together confirm the extensive adaptive response induced by 20S proteasome hyperactivation. Additionally, the enrichment on the chaperone subnetwork revealed significant enrichment on small HSPs, oxidative stress defense factors, and Hsp70 family of proteins (Fig. 2G-H; Hypergeometric distribution FDR<0.05), pathways widely recognized for their role in maintaining proteostasis, promoting longevity, and enhancing stress resilience in *C. elegans*. These findings highlight the selective involvement of key proteostasis components in the adaptive response, pointing to alterations in the proteostasis network due to 20S proteasome hyperactivation. Within the oxidative stress defense, SOD-2 and SOD-3 were elevated, while cytoplasmic SOD-1 and gpx-5 declined (Table S1), suggesting a reduced need for cytoplasmic defenses in α3ΔN. Concurrently, several sHSPs were downregulated (Table S1), indicating that 20S gate opening diminishes reliance on conventional proteostasis machinery. We next further tested this hypothesis.

### Proteasome hyperactivation enhances oxidative stress defense independently of SODs enzymes

With the observed reduction in cytosolic oxidative response factors in α3ΔN we hypothesized that proteasome hyperactivation may bolster oxidative stress resistance. To assess whether oxidative damage is reduced in the α3ΔN strain, we analyzed post-translational modifications (PTMs) focusing on oxidation-induced modifications in our Tandem Mass Tag (TMT) dataset. We identified 1,997 peptides exhibiting signs of oxidative modifications. Among these, 63 peptides showed a significant decrease in oxidation levels, while 22 were significantly upregulated (Fig. 3A-TableS4). The reduction in oxidative markers can be explained by increased degradation of oxidatively damaged proteins. This is consistent with the finding that the 20S proteasome is capable of degrading oxidatively damaged IDPs (42), which would be enhanced by 20S gate-opening. An additional consideration is that basal oxidative stress in the α3ΔN mutants maybe be reduced. These results indicate a more robust and less damaged proteome, particularly in aging worms, which may confer increased resistance against oxidative stress, which we directly test below.

**Figure 3:**
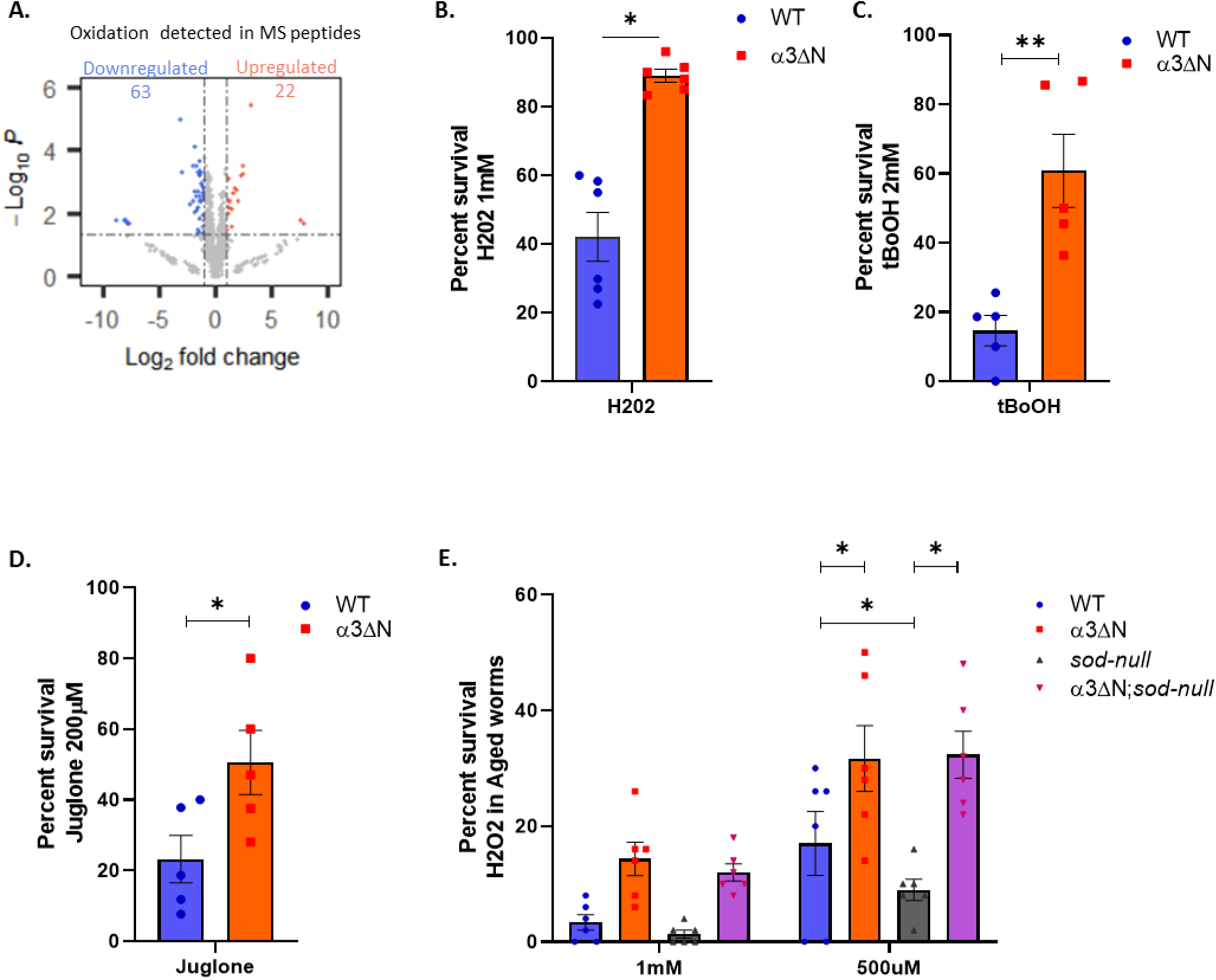
Enhanced oxidative stress resistance in α3ΔN mutants is independent from SODs. **A.** Volcano plot of the differentially post-translation modified (PTM) oxidated peptides detected in the TMT-MS with Maxquant *(Q<0.01).* Blue points above the x axis dotted line represent oxidated peptides downregulated (FC<-1.5) and yellow dots represent oxidated peptides upregulated (FC>1.5). Distribution of peptides was adjusted with Limma model and significance was determined with ANOVA and Tukey correction for multiple test*(FDR<0.05)*. **B-D.** Percentage of surviving of WT and α3ΔN, 1 day adult worms after acute oxidative stress challenge with H2O2 at 1mM (B), tBoOH at 2mM (C) and juglone at 200μM (D). Data represents mean values ± SEM of at least four independent experiments \**p* < 0.05, \*\**p* < 0.01 by unpaired two-tailed *t*-test. Total number of worms assessed for H2O2: (α3ΔN: 335 ; WT:351); tBoOH(α3ΔN: 175 ; WT:171); Juglone (α3ΔN: 389 ; WT:394). **E.** Percentage of surviving old age worms (Day 8) of the indicated genotypes after acute oxidative stress challenge with H2O2 at 1mM and 500 μM. *sod-null*, knockout worms for SODs and α3ΔN;*sod-null* indicates open gate worms crossed into *sod-null* knockout strain. Data represents mean values ± SEM of five independent experiments \**p* < 0.05 by two-way-ANOVA with Sidaks correction for multiple comparisons-test. Total number of worms tested at 1mM (α3ΔN: 520 ; WT:468; *sod-null* : 600; α3ΔN; *sod-null*: 500), at 500uM (α3ΔN: 600 ; WT:600; *sod-null*: 400; α3ΔN; *sod-null*: 400).

To confirm if proteasome hyperactivation contributes to oxidative stress defense, we subjected α3ΔN mutants to acute oxidative stress using various compounds that induce reactive oxygen species (ROS) in the cytosol and mitochondria. Hydrogen peroxide (H₂O₂) is particularly relevant for *C. elegans*, as it is produced by pathogenic bacteria in their natural environment (50, 51). We also used tert-butyl hydroperoxide (tBoOH), a stable cytosolic ROS generator (52), and juglone, a mitochondrial stressor known to induce ROS through activation of the FLP neuropeptide family (53). Aldicarb, an acetylcholinesterase inhibitor used for assessing synaptic transmission (54), served as a non-ROS control stressor.

Our results revealed that α3ΔN mutants displayed markedly enhanced resistance to the oxidative stressors H₂O₂, tBoOH, and juglone (Fig. 3B-D), while their response to the neurotoxic agent aldicarb remained unchanged (as it functions via a non-ROS mechanism) (Fig. S4A), confirming that the observed resistance is specific to oxidative stress. Probit curve analysis quantified the lethal dose 50 (LD₅₀) and resistance ratios (RR) for each compound (Fig. S4B-E). The analysis indicated that a 3.43-fold higher concentration of H₂O₂ was required to affect 50% of the α3ΔN mutant population (Fig. S4B,E). Similarly, the mutants required a 4.25-fold increase in tBoOH concentration (Fig. S4C,E) and a 1.84-fold increase for juglone to reach the LD₅₀ threshold (Fig. S5D,E). These results demonstrate that α3ΔN is highly resistant to multiple forms or ROS stress, but what if we remove natural cellular ROS defenses, can the α3ΔN still protect against ROS?

SODs are well-known key effectors of oxidative stress defense, as they catalyze the dismutation of superoxide radicals into oxygen and hydrogen peroxide, thereby mitigating oxidative damage. To understand the role of SODs in the observed resistance, we crossed the α3ΔN mutants with the MQ1766 strain, which lacks all SOD genes (hereafter referred as sod-null) and is known to be sensitive to oxidative stressors (55). Previous studies have demonstrated that deleting even a single major sod gene, such as sod-1 or sod-2, is sufficient to markedly increase sensitivity to oxidative stress in *C. elegans* (55, 56), and thus the sod-null is extremely sensitive to oxidative stress. We crossed this sod-null with α3ΔN. While the mitochondrial sod-2 could not be crossed out, due to its close proximity to pas-3 (2 centiMorgans apart) all other sod genes were knocked out resulting a quadruple sod mutant in the α3ΔN background. Intriguingly, although the sod-null mutation diminished stress resistance overall relative to wild-type, the α3ΔN sod-null mutants retained significant resistance to tBoOH and H₂O₂ stress, even into advanced age (Fig. 3E, Fig. S4E). In fact, the sod-null worms with the hyperactive proteasome showed no difference in H2O2 sensitivity at 500μM compared to α3ΔN. The exception was juglone, which exhibited similar toxicity in sod-null worms regardless of the α3ΔN mutation, likely because juglone induces mitochondrial ROS and mitochondria lack proteasomes. Although SOD enzymes serve as primary defenses against oxidative stress by neutralizing superoxide radicals, the substantial resistance to oxidative stress observed in α3ΔN mutants, despite the absence of most SODs, demonstrates the substantial protective capacity of the 20S proteasome. In addition, the fact that the enhancement of IDP degradation is highly protective during ROS stress suggests that IDP’s may be a particularly vulnerable protein species in oxidative tissues like the brain.

Our findings suggest that instead of relying on enzymatic detoxification of ROS, the hyperactive proteasome enhances the clearance of oxidatively damaged proteins, thereby reducing the accumulation of damaged macromolecules and maintaining proteome integrity. This represents a novel mechanism of oxidative damage mitigation, strengthening cellular defenses through improved protein degradation rather than traditional antioxidant pathways. The consistent resistance of α3ΔN mutants to H₂O₂ in older worms indicates the presence of a durable system capable of handling protein damage throughout the lifespan of *C. elegans* (Fig. 3E). These findings provide mechanistic insight into how proteasome hyperactivation fortifies organisms against oxidative damage by bolstering stress response capacity, enhancing resilience to various stressors even independently of SOD enzymes.

### Elevated ERAD capacity through proteasome hyperactivation mitigates vitellogenin aggregation

While our data above demonstrates that hyperactivation of the 20S proteasome could accelerate the degradation of IDPs and oxidatively damaged proteins we did not expect the α3ΔN-20S to have impacts on ERAD since it is well known that the 26S proteasome mediates ERAD after P97/VCP extracts substrates from the ER. To determine if ERAD was affected we treated both wild-type and α3ΔN worms with tunicamycin to assess whether proteasome hyperactivation could impact ER stress response. Tunicamycin induces ER stress by inhibiting N-linked glycosylation leading to protein misfolding in the ER (57). We were surprised to find that α3ΔN mutants were significantly protected from tunicamycin compared to wild-type worms (Fig. 4A). These results indicate 20S hyperactivation protects against ER misfolding stress. Therefore, we anticipated that some aberrant, ER resident aggregation-prone proteins could be selectively targeted by the hyperactive proteasome. Indeed, our proteomic analysis revealed that the entire vitellogenin family, key proteins associated with aging in *C. elegans*, were significantly downregulated at the protein level (Fig.S3C), while RNA-seq data showed no significant changes at the transcript level (Fig.S3D). This suggests that proteasome hyperactivation promotes the degradation of vitellogenins post-translationally, rather than affecting their transcription. In normal aging of *C. elegans*, dysregulation of vitellogenin expression leads to the abnormal accumulation of yolk proteins and lipoproteins, resulting in adverse effects such as senescent proteinopathy, organ degeneration, obesity, and proteotoxicity (58–61). Interestingly, vitellogenins in *C. elegans* are homologous to human apolipoprotein B (ApoB) (62), which is degraded via the ERAD pathway (63), hence we expect that the proteasome also catalyzes vitellogenin degradation. In doing so, these lipoproteins become central targets through which proteasome hyperactivation improves proteostasis by preventing their usual buildup into damaging aggregates.

**Figure 4.**
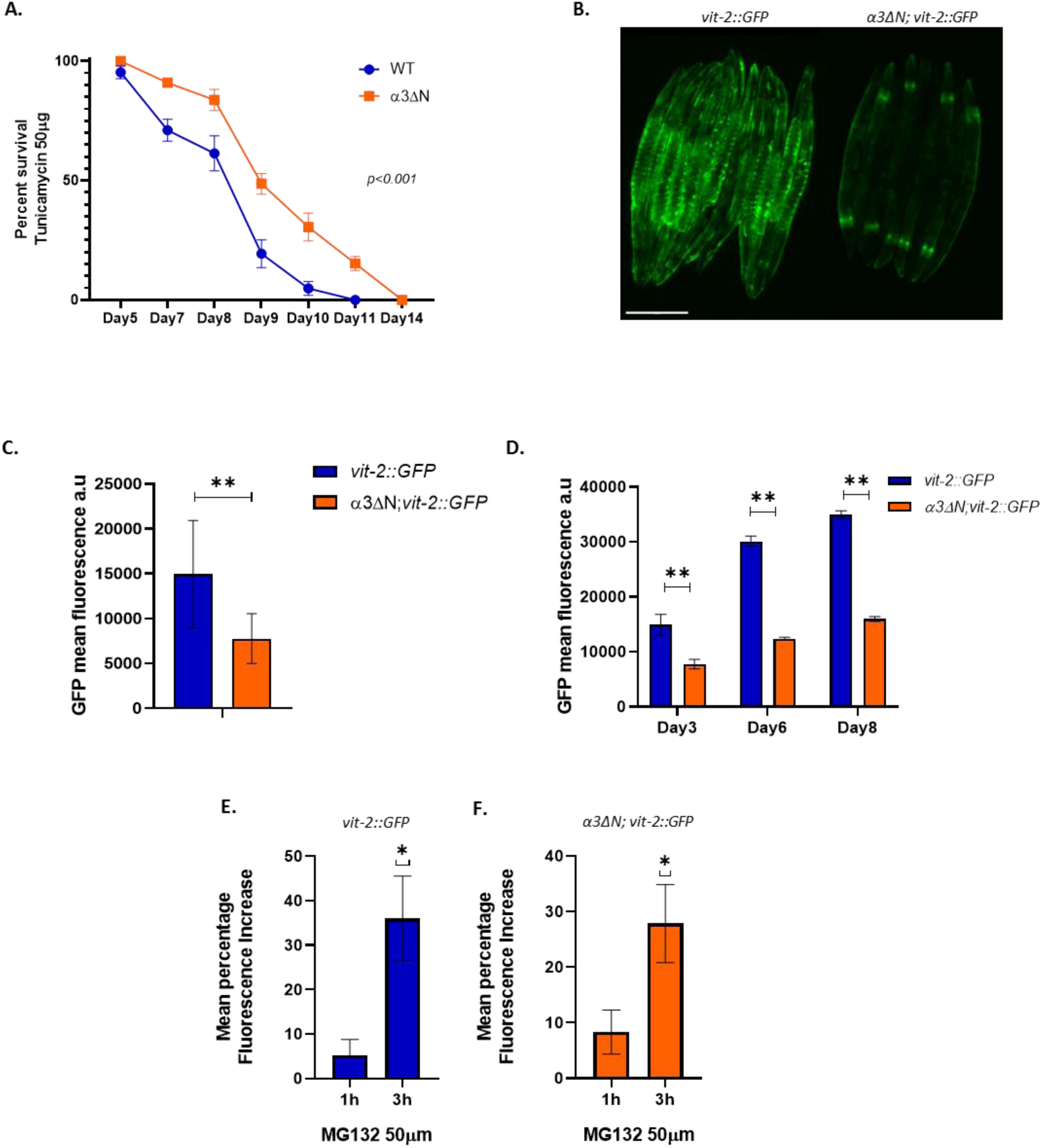
Enhanced ERAD in α3ΔN mutants reduces vitellogenin accumulation. **A.** Survival of WT animals and α3ΔN Mutants upon tunicamycin treatment 50 (μg/mL) (data from *n* = 3 independent experiments) *p<0.01* determined by the log-rank test. Representative images **(B**) and fluorescent quantification (**C**) of *vit-2::GFP + rol-6(su1006)* and α3ΔN;*vit-2::GFP + rol-6(su1006)* worms expressing the vitellogenin 2 protein fused to GFP vit-2::GFP from the outcrossed RT99 strain with defective accumulation of vit-2. Scale bar 500μM. Images of day 3 worms at 4x magnification. Data in **C** represents mean values ± SD \*\**p* < 0.01 by two-tailed *t*-test. **D.** *vit-2::GFP* Accumulation Over Time. We tracked the fluorescent intensity of *vit-2::GFP* versus α3ΔN; *vit-2::GFP* worms, revealing decrease in *vit-2::GFP* accumulation in α3ΔN;*vit-2::GFP* mutants. Statistical analysis *(**p < 0.01*) via multiple two-tailed t-tests. Data shown as mean ± SD. **E-F.** *vit-2::GFP* validation as proteasome substrate by quantifying GFP fluorescence after addition of proteasome inhibitor MG132 (50μM) to the *vit-2::GFP + rol-6(su1006)* (E) and α3ΔN; *vit-2::GFP + rol-6(su1006)* (F) adult worms. Data represents mean values ± SD (n=6) \**p* < 0.05 by two-tailed *t*-test with welch correction.

We proposed that by opening the 20S proteasome gate via the α3ΔN-20S mutation could enhance degradation by the ERAD pathway. This hypothesis is mechanistically compelling because ERAD substrates are unfolded by P97/VCP to extract them from the ER and release them into the cytosol, making them potential substrates for the α3ΔN-20S proteasome. Based on this mechanism, we predicted that these mutants would show accelerated degradation of specific ER substrates explaining the MS results above. To validate this, we specifically monitored vitellogenin levels, by using a *C. elegans* strain expressing a Vitellogenin-GFP fusion protein (vit-2::GFP) from the RT99 strain with no rme-4 mutation (Fig4B). After crossing our α3ΔN mutants with our vit-2::GFP strain, we quantified GFP levels. The results showed a significant decrease in vit-2::GFP fluorescence in the α3ΔN worms (50% reduction of vit-2::GFP consistently across the population) (Fig 4B-C) and this decline remained consistent through the aging process (Fig4D), corroborating the vitellogenin depletion detected in our proteomic data and supporting enhanced ERAD in the α3ΔN mutants. To confirm the reduction in vit-2::GFP was due to proteasome degradation we also inhibited proteasome activity with MG132. Within three hours of exposure, we observed a significant increase in GFP fluorescence in WT and α3ΔN worms treated with MG132 (Fig4E-F). This increase in fluorescence directly supports the hypothesis that vit-2::GFP protein levels are regulated by proteasomal degradation, which is enhanced by proteasome hyperactivation. In addition, this data supports the hypothesis that vitellogenins are key targets through which the 20S proteasome improves proteostasis in the hyperactive state.

### Proteasome hyperactivation enhances the clearance of pathogenic human ATZ protein

The significant reduction of vitellogenins via enhanced ERAD in α3ΔN mutants prompted us to investigate whether proteasome hyperactivation could be effective against other misfolded ER proteins implicated in human diseases. So, in an orthogonal approach we crossed the α3ΔN into a *C. elegans* model for Alpha-1 antitrypsin deficiency (A1ATD), a condition marked by accumulation of ATZ in the ER (64). Due to a genetic mutation of ATZ in humans, these misfolded proteins accumulate in the liver, hindering its function and elevating the risk of liver diseases (37, 65). ATZ is directly associated with the ERAD pathway, making it heavily reliant on proteasome activity for the clearance of misfolded alpha-1 antitrypsin proteins. Remarkably, α3ΔN mutants showed significantly reduced accumulation of sGFP-ATZ (Fig5A-B), and this reduction persisted with aging (Fig5C-D), indicating improved degradation and clearance of these aberrant proteins. In addition, to determine that clearance of ATZ aggregates was due to proteasome activity we again used the proteasome inhibitor MG132 on both the ATZ-expressing worms and those crossed with the α3ΔN mutant. We observed a significant increase in the number of ATZ aggregates in both strains after proteasome inhibition (Fig5F-G). This demonstrates that the efficient degradation of ATZ aggregates is indeed reliant on proteasome function in the α3ΔN strain. In addition, these findings demonstrate that the α3ΔN-20S proteasome can plays a role in ER-associated degradation pathways and maintains protein homeostasis during stress conditions triggered by protein aggregation. Moreover, this work establishes α3ΔN-20S proteasome activation as a promising therapeutic strategy for treating disorders like ATZ, where protein homeostasis mechanisms are compromised.

### Proteasome hyperactivation increases ER stress resistance and lifespan independently of the Unfolded Protein Response

Because enhanced ERAD in α3ΔN appears to play a role in the extensive downregulation of vit-2 in the ER (Fig4F), which could also affect other ER resident proteins, we sought to further determine if 20S hyperactivation may affect the cytosolic UPR, ER’s UPR or the mitochondrial UPR (mtUPR). To do so, we crossed the α3ΔN strain with three chaperone reporter strains. The strain CL2070, which expresses hsp-16.2p::GFP (dvIs70), a small heat shock protein widely used as a biomarker for cytosolic heat stress (66). We selected this reporter based on our enrichment analysis indicating changes in sHSP. The strain SJ4005, expressing hsp-4::GFP (zcIs4), which serves as a marker for UPR activation due to its homology to the ER chaperone BiP (67) and GL347, expressing hsp-6p::GFP (zcIs13; along with lin-15(+)), which acts as a reporter of mtUPR induction (68). . These strains were crossed with α3ΔN and basal UPR responses were quantified. These results revealed no significant changes in mtUPR activation in α3ΔN mutants compared to wild-type worms through the lifespan, as measured by hsp-6p::GFP fluorescence (Fig. S5A-B). This demonstrates that mitochondrial stress is not affected by proteasome hyperactivation and thus should not contribute to the observed phenotypes (e.g. ER stress resistance or longevity). Notably, under normal conditions we observed no induction of hsp-16.2 mirroring our findings for the mtUPR reporter (Fig. S5C-D). Together with the lack of hsp-16.2 induction, these results suggest that under normal conditions, neither cytosolic heat shock nor mitochondrial stress pathways are activated, emphasizing that the selective enhancement of ERAD is the primary mechanism by which 20S hyperactivation targets misfolded and aggregation-prone proteins. However, α3ΔN mutants crossed with hsp-4::GFP, the UPR reporter, exhibited increased fluorescence under normal conditions, suggesting elevated expression of hsp-4 and potential ER UPR activation.

Moreover, the reporter activity started very early in the life of our α3ΔN mutants, from the larval stage L1 and this increased hsp-4::GFP activity persisted into late life stages (Fig.6A-B). This sustained increase in hsp-4 levels in the α3ΔN mutants could contribute to α3ΔN’s tunicamycin resistance phenotype and increased lifespan. Normally, hsp-4 is induced by the UPR-ER pathway and is dependent on xbp-1, which acts as an important regulator of stress resistance and longevity in *C. elegans* (69). To assess whether the induction of UPR is necessary for the increase tunicamycin resistance and lifespan extension in α3ΔN mutants, we introduced an xbp-1 loss of function xbp-1αΔαΔ—(zc12) mutation into the α3ΔN;hsp-4::GFP background. This mutation effectively disrupted the IRE-1/XBP-1 branch of the UPR in the xbp1 crosses, as expected, since tunicamycin-mediated induction of hsp-4 was abolished (Fig. 6C-D). Remarkably, the α3ΔN; hsp-4::GFP; xbp-1 null triple mutants maintained substantial resistance to tunicamycin compared to xbp-1 null mutants alone, even remaining similar to wild-type worms (Fig. 6E). This significant resistance, even in the absence of xbp1 activation and BiP induction, suggests that the hyperactive proteasome in α3ΔN effectively compensates by robustly clearing misfolded proteins, thereby alleviating ER stress in the absence of this key protective pathway. Moreover, we further assessed the lifespan of these xbp1 crosses to determine if xbp1 activation was necessary for α3ΔN extended longevity. We were surprised to find that the worms still maintained the extended lifespan phenotype (Fig.6F), indicating that longevity conferred by proteasome hyperactivation is independent of UPR signaling via the XBP-1 pathway. Taken together these results indicate that the enhanced 20S proteasome can independently function as a protective proteostasis pathway that has capacity to reduce proteotoxic stress and increase longevity.

## Discussion

The accumulation of misfolded and aggregated proteins, particularly intrinsically disordered proteins (IDPs), is a hallmark of Aging-related proteinopathies and neurodegenerative conditions, implicating the role of the proteostasis network and the proteasome system in maintaining cellular health (15, 67, 68). In this study, we demonstrate that constitutive activation of the 20S proteasome achieved via the α3ΔN mutation provokes a dramatic remodeling of both the proteome and transcriptome in *C. elegans*. Approximately 10% of the proteome is altered, accompanied by a network-wide shift in transcription activity. This global reprogramming is paralleled by a coordinated increase in both protein synthesis and degradation rates (Fig 1-2). This adaptive coordination demonstrates the cells remarkable resilience to adapt to increased degradation rates by accelerating protein synthesis. Importantly, our heat treatment method, which enriches for aggregation-prone proteins, reveals a 50% reduction in heat-stable proteins (e.g. IDPs) in the α3ΔN strain (Fig 1). Similarly, Pepelnjak, M. et al identified that many 20S substrates are RNA and DNA-binding proteins with intrinsically disordered regions (42). Moreover, tau degradation was also reported in a α3ΔN-20S overexpression cell model (70), consistent with these results. In addition, organismal studies employing proteasome overexpression have reported increased stress resistance and extended lifespan, further corroborating our results (23–27). However, our approach fundamentally differs in that rather than overproducing proteasome components, we induce a structural change that constitutively opens the 20S proteasome gate. This structural modification triggers the phenotypes we observe without altering total proteasome expression levels. Because neurodegenerative diseases are driven by the accumulation of IDPs like tau, α-synuclein, TDP-43, and huntingtin, as well as many others, this 20S degradation pathways becomes highly relevant to these diseases. Therefore, our novel finding directly links the open 20S proteasome configuration to the preferential clearance of misfolded and IDPs substrates critical in the pathogenesis of neurodegeneration. substrates critical in the pathogenesis of neurodegeneration.

Additionally, our data show that hyperactivation of the 20S proteasome enhances resistance to oxidative stress, which is associated with aging and various neurodegenerative diseases. We observed reduced levels of oxidative damage markers in TMT-MS peptides. In addition, we found that α3ΔN was resistant to ROS-generating agents such as H₂O₂ and tert-butyl hydroperoxide. Surprisingly, the open 20S was even protective in a SOD-deficient background (Fig 3). This challenges the paradigm that ROS detoxification relies solely on enzymatic antioxidants. Instead, the 20S proteasome appears to be a strong contributor to the mitigation of oxidative damage by enhancing the clearance of oxidatively damaged proteins. This SOD-independent ROS resilience suggests that coupling proteasome activation with traditional antioxidant pathways could amplify cellular defenses in aging and disease. These results also indicate that the enhanced degradation of oxidatively damaged IDP’s is a likely contributor to the longevity phenotype observed in the α3ΔN worms. However, the α3ΔN mutants also exhibits significant resistance to tunicamycin induced ER stress, likely due to enhanced ER-associated degradation (ERAD) that was observed. This stress resilience in the cyto/nuclear and ER compartments reveals novel and integrated benefits of 20S proteasome hyperactivation combining into a robust platform for proteotoxic stress resistance, which is critical determinant for health organismal aging.

A major novel finding of our work is the enhanced capacity of ERAD in the α3ΔN mutants. ERAD is essential for clearing misfolded proteins from the endoplasmic reticulum, and its failure contributes to many diseases. Vitellogenins, naturally processed by the ER and prone to aggregation, serve as endogenous substrates whose downregulation in our model provides direct evidence of enhanced ER degradation and mirrors the proteasome mediated clearance of ApoB in mammals, highlighting its conserved role in lipoprotein homeostasis. The robust depletion of vitellogenins, together with the observed tunicamycin resistance (Fig 4) prompted us to evaluate additional ERAD substrates. Indeed, the marked reduction in ER resident misfolded human ATZ protein levels further emphasizes that the constitutively open 20S proteasome efficiently accelerates the degradation of even aggregation-prone proteins originating from the ER (Fig 5). Therefore, these results position 20S hyperactivation as a viable strategy for disorders marked by ERAD dysfunction. The finding that the α3ΔN-20S could accelerate ERAD function was surprising since the 26S is a known endpoint for ERAD substrates, which are typically ubiquitinated after extraction. While the α3ΔN mutation does slightly enhance the 26S dependent degradation of a linear chain Ub4-GFP substrate, peptide degradation rates, which are gating dependent, are similar for 26S and α3ΔN-26S complexes via in-gel activity assay (28). Since P97 must unfold ER resident proteins to extract them into the cytosol compartment, it seems more likely that the α3ΔN-20S, which is activated >100X more than the WT 20S (28), is responsible for the enhancement of ERAD that we observe in α3ΔN animals.

**Figure 5.**
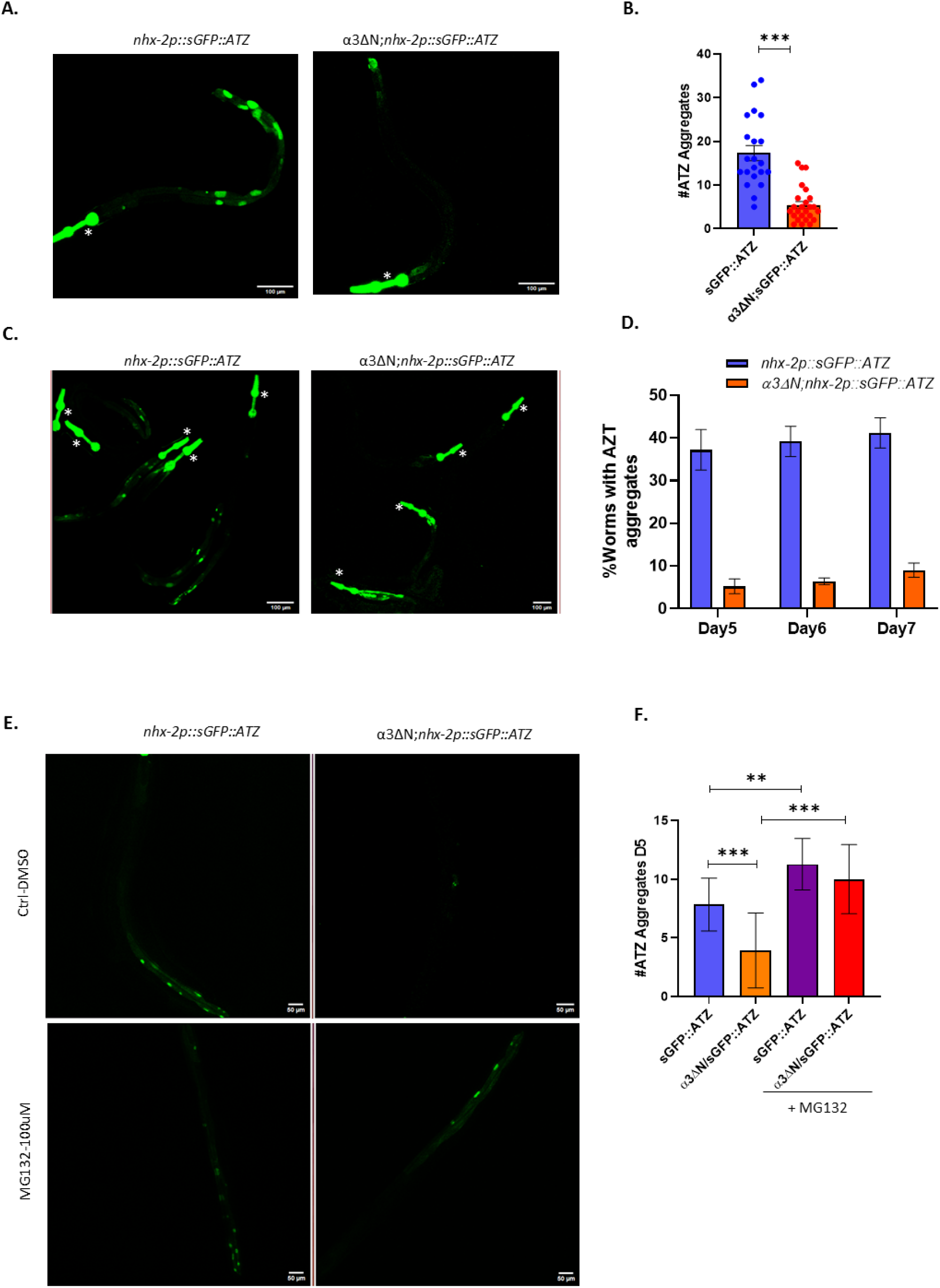
Enhanced ERAD in α3ΔN mutants reduces pathogenic ATZ accumulation. **A.** Representative Imaging of day 7 transgenic animals expressing ERAD substrate *nhx-2p::sGFP::ATZ* in intestinal epithelium. Scale bar, 100 μm. **B.** ATZ Aggregate Quantification: Comparison of ATZ aggregate counts in *nhx-2p::sGFP::ATZ* (n=21) and α3ΔN; *nhx-2p::sGFP::ATZ* (n=25) worms on day 7. Data represents mean values ± SEM \*\*\**p* < 0.001 by two-tailed unpaired *t*-test**. C.** Day 5 worms expressing ATZ ERAD substrate. Visualization in gut epithelium of transgenic worms expressing GFP-tagged ERAD substrates (sGFP::ATZ) *nhx-2p::sGFP::ATZ* compared to α3ΔN; *nhx-2p::sGFP::ATZ* .Scale bar 100 μm. The bright pharyngeal signal, indicated with an asterisk *, is a genetic marker and should not be mistaken for ATZ aggregates. **D.** ATZ Aggregate accumulation over time in α3ΔN Mutants. Tracking down the formation of ATZ aggregates demonstrates a reduction in ATZ aggregates in α3ΔN; *nhx-2p::sGFP::ATZ* mutants compared to *nhx-2p::sGFP::ATZ* across adulthood, with n=50 for both groups at specified days, showing the enhanced degradation capabilities. **E.** Representative images of *nhx-2p::sGFP::ATZ at day 7* after treatment with the proteasome inhibitor MG132 (100 μM). Images show increased accumulation of ATZ aggregates in both *nhx-2p::sGFP::ATZ* and α3ΔN; *nhx-2p::sGFP::ATZ* worms upon proteasome inhibition. Scale bar, 100 μm. **F.** Quantification of ATZ aggregates from (E). Data represent mean values ± SEM (n = at least 15 worms per group). p < 0.01 determined by one-way ANOVA followed by Tukey’s post hoc test.

Mechanistically, our findings also challenge the traditional view of the 20S proteasome as merely a passive core within the 26S complex. Instead, our data indicate that an open 20S proteasome functions as a central regulator of proteostasis by directly targeting IDPs and unfolded proteins. Notably, our genetic crosses with proteostasis reporter strains revealed that mitochondrial and cytosolic stress pathways are unperturbed, though we do see induction the ER-specific hsp-4 reporter from early life stages. This selective UPR activation could explain the enhanced resistance to ER stressor tunicamycin. This observation is important because UPR activation has been shown to underlie many long-lived models in *C. elegans* (69, 71, 72) aand is an important pathway in many proteinopathies. We fully anticipated that the hsp-4 upregulation was driving the longevity phenotype in α3ΔN, but its cross with mutant xbp1 stain proved otherwise. Surprisingly, these results showed that ER stress resistance and increased lifespan in the α3ΔN strain were still retained even in the absence of xbp1 mediated UPR activation (Fig 6). These findings indicate that the enhancement of IDP degradation and ERAD function conferred by inducing 20S gate-opening is the primary mechanism driving the proteotoxic stress resistance and longevity phenotypes observed in the α3ΔN strain. This suggests that the 20S proteasome is an important mediator of cellular proteostasis. This hypothesis could be further supported by approaches that impair 20S function without affecting 26S function, but major technical hurdles must be overcome first to enable such capabilities.

**Figure 6.**
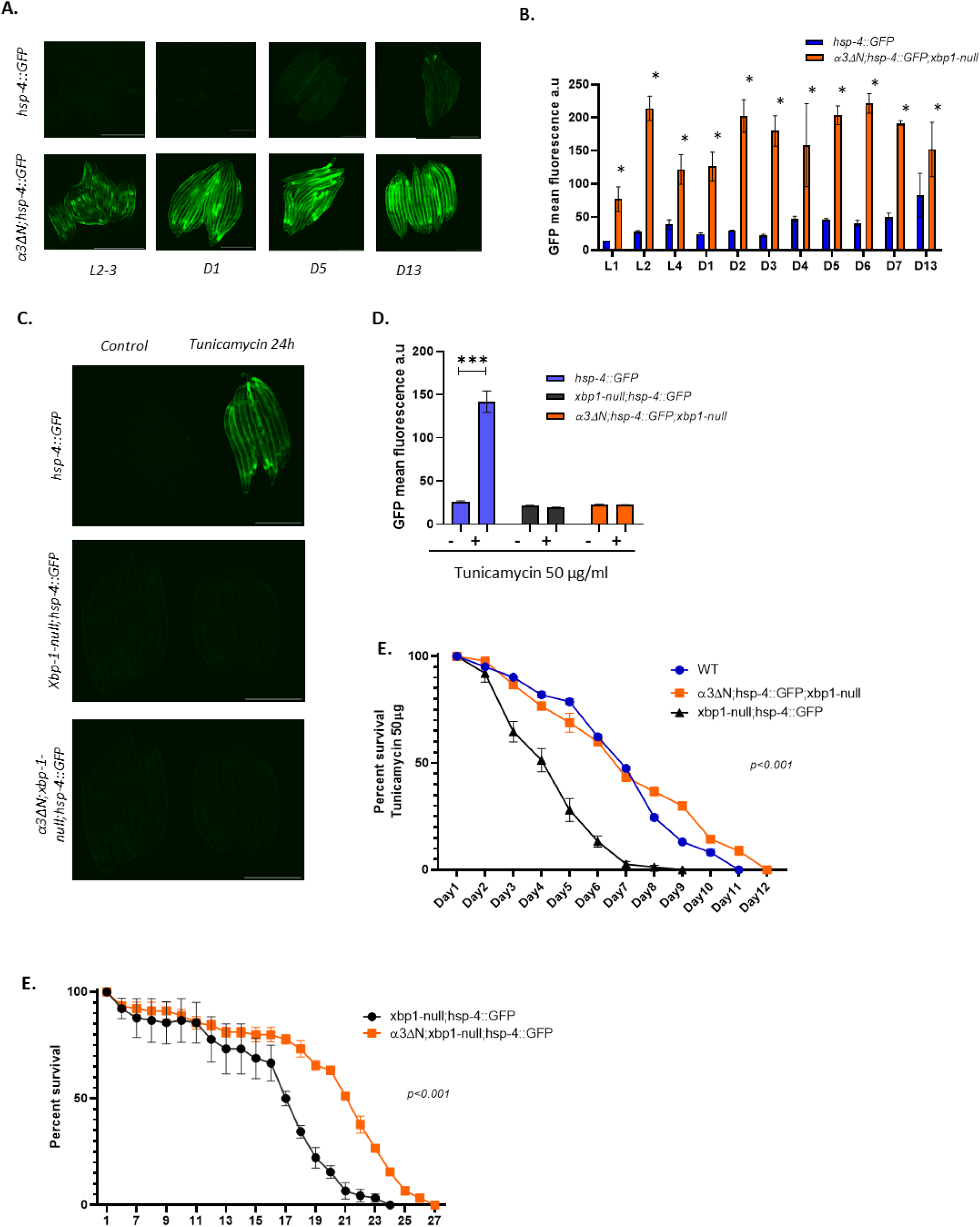
Enhanced ERAD and extended lifespan in α3ΔN mutants occur independently of UPR activation via hsp-4 (BiP) and xbp-1. **A** Representative time-course images of hsp-4::GFP and α3ΔN;hsp-4::GFP expression starting at the 24-hour larval stage (L1). Images were captured at 20× magnification. Scale bar, 500 μm. **B** Quantitative fluorescence analysis of hsp-4::GFP expression over time in hsp-4::GFP and α3ΔN;hsp-4::GFP worms. Data are presented as mean ± SD from at least three independent experiments *(*p < 0.01*, two-way ANOVA). **C.** Representative images of ER UPR activation in day 1 adult worms following treatment with either 50 μg/ml tunicamycin or control DMSO for 24 h at 20°C. Shown are hsp-4p::GFP, hsp-4p::GFP; xbp-1(zc12) (xbp-1 null), and triple mutant α3ΔN;hsp-4p::GFP; xbp-1(zc12) worms. Scale bar, 500 μm. **D.** Quantification of GFP fluorescence intensity from Panel **C**. In hsp-4p::GFP control strains, tunicamycin induced a robust increase in hsp-4 (BiP) expression, whereas α3ΔN;hsp-4p::GFP;xbp-1(zc12) mutants showed no significant induction, confirming the abolition of tunicamycin-mediated BiP upregulation. Data are presented as mean ± SEM (*p < 0.01, two-way ANOVA). **E.** Survival analysis under tunicamycin stress. At day 1 of adulthood, WT (blue), xbp-1(zc12) null (black), and α3ΔN;hsp-4p::GFP; xbp-1(zc12) worms were transferred to plates containing 50 μg/ml tunicamycin, and survival was monitored. Median lifespans were 7 days for WT, 5 days for xbp-1(zc12) null, and 7 days for α3ΔN;hsp-4p::GFP; xbp-1(zc12) (*p < 0.0001*). **F.** Kaplan–Meier survival curves for hsp-4p::GFP; xbp-1(zc12) (N = 90) and α3ΔN;hsp-4p::GFP; xbp-1(zc12) (N = 90) worms maintained on FuDR plates from day 1 of adulthood. Median lifespans were 17.5 days for hsp-4p::GFP; xbp-1(zc12) and 22.0 days for α3ΔN;hsp-4p::GFP; xbp-1(zc12) (*p < 0.0001*).

Surprisingly, constitutive proteasome hyperactivation in *C. elegans* has limited downsides. One potential trade-offs is the reduction in fecundity, but this concern is irrelevant in therapeutic context and no other negative phenotypes have been observed in the α3ΔN mutants. It is also important to clarify that the eating defects found in our model, which show approximately a 10% decrease (28), are minor compared to established longevity mutants like eat-2 mutants, which exhibit about a 90% reduction in feeding (73, 74). Importantly, the worms appear nominal and have normal motor control, indicating no perturbations to neuronal or developmental functions. In addition, WT and α3ΔN both respond similarly to aldicarb, a neurotoxin indicating that neurons in the α3ΔN function normally and are not stressed. Therefore, therapeutic manipulation of 20S proteasome activity appears to offer distinct advantages over current strategies. 20S activators could selectively enhance degradation of toxic IDPs and oxidatively damaged proteins while sparing folded substrates. This is particularly advantageous since IDPs are the primary protein class associated with neurodegenerative disease, and are particularly sensitive to oxidative damage since all of their residues are exposed to ROS, unlike folded proteins which contain many buried and protected residues. In addition, pharmacological targeting of the 20S gate is mechanistically feasible since peptides mimicking the HbYX motif or small peptide mimetics like ZYA are able to robustly induce 20S gate opening (20).

In summary this work redefines 20S proteasome function as an important pathway in cellular health and as a tunable tool capable of mitigating age-related decline through IDP degradation, oxidative damage clearance, and ERAD enhancement. Its independent function distinct from canonical stress responses highlights its potential as a therapeutic modality for neurodegenerative diseases, alpha-1 antitrypsin deficiency, and other proteinopathies. Future studies should map structural determinants of 20S substrates (e.g., disorder, hydrophobicity) that may regulate 20S function and explore combinatorial approaches with autophagy modulation to optimize proteostasis resilience.

## Materials and Methods

### C. elegans strains and culture

The open gate mutant strain pas-3(dsm100) was utilized to study proteasome hyperactivation effects. Throughout this study, this strain is referred to as α3ΔN. C. elegans strains were cultured on nematode growth medium (NGM) agar plates at 20°C, with E. coli strain OP50 serving as the food source, following standard protocols (74). The N2 Bristol strain was used as the wild-type (WT) control. Possible confounding effects due to differences in fertility were controlled for using FUDR (a standard sterilizing agent) in all the experiments unless stated otherwise. Therefore, strains were under the same fertility conditions. In addition, unless stated otherwise, all experiments were conducted on day 1 of adulthood, focusing exclusively on hermaphrodite worms. Detailed information on the strains utilized in this study can be found in Supplementary Table S5. Age synchronization was performed via alkaline bleaching. Gravid adults from two 10-cm plates were collected and washed three times with ddH₂O in 15-mL conical tubes. After reducing the volume to 3.5 mL, a 1.5 mL solution of bleach/NaOH (1 mL 5% sodium hypochlorite and 0.5 mL 5N NaOH) was added. The mixture was vortexed every 2 minutes for 10 minutes until no nematode fragments remained. Sterile M9 buffer was then added to neutralize the reaction, and the tubes were centrifuged at 1,100×g for 1 minute to pellet the eggs. Eggs were washed with 10 mL sterile M9, centrifuged at 2,000×g for 1 minute, and the supernatant discarded. The pellet was resuspended in 7.5 mL sterile M9 in a fresh 15-mL tube and incubated overnight with gentle agitation to allow hatching. L1-arrested nematodes were harvested within 24 hours, plated on OP50-seeded plates, and incubated at 20°C until the desired stage.

### TMT-MS-Proteomic changes in α3ΔN mutants

Worms were synchronized, then populations grown on plates with FUDR. Protein extracts from 8-day old adult populations, were processed using a CME/FASP Digest for trypsinization, followed by labeling with TMT10plex Isobaric Tagging for multiplex quantitative analysis. Post-labeling, samples were fractionated via bHPLC, employing a linear gradient from 5% to 35% acetonitrile in water over 60 minutes, with a flow rate of 300 µL/min, using a C18 column (20 cm length, 1.7 µm particle size). Fractions were collected every 2 minutes. These were then analyzed on an Orbitrap Eclipse mass spectrometer for high-resolution detection using TMT MS3; 60 min gradient per fraction. Data were then processed using MaxQuant (75) against the C. elegans protein database (75) to identify proteins and extract MS3 reporter ion intensities, with search parameters including trypsin specificity, mass tolerances of ±10 ppm for precursors and ±0.02 Da for fragments, fixed modifications for TMT tags on lysine and peptide N-termini and carbamidomethyl on cysteine, variable modifications for oxidation on methionine, peptide and protein FDR thresholds set at 1%, and quantification based on MS3 reporter ion intensities. Proteins identified as decoys and contaminants were filtered out, as well as rows with more than 80% zero values (count = 0). To handle zero values and facilitate logarithmic transformation, a constant value of +1 was added to each intensity value in the filtered dataset (76–79). The adjusted dataset was then loaded into a matrix of intensities and processed in R version 3.6.3 for log₂ ratio conversion using the median sweeping method for normalization. After normalization, Principal Component Analysis (PCA) was plotted to visualize the expression patterns among the samples. For Differential Expressed Proteins quantification DEqMS package was used (40) since this method consider the specific structure of MS data and fitted best our data compare to others widely used such as Limma (80). Proteins with fold changes > |1,5| and false discovery rates (FDRs) < 0.05 were defined as Differential Expressed Proteins (DEPs) and captured for functional enrichment analysis. Regarding detection and quantification of post-translational modifications (PTMs), focusing on oxidation in α3ΔN mutants we employed MaxQuant which facilitated the identification of proteins and their oxidation states by matching mass spectra against the C. elegans protein database as explained above, utilizing a site-level false discovery rate (FDR<00.1) to ensure accurate PTM identification (75). Quantitative data extracted from MaxQuant, including modification-specific peptides’ intensities and ratios, were then analyzed to assess the extent of oxidation in R under the Limma package (40).

### RNA-seq Transcriptomic changes in α3ΔN mutants

Total RNA was extracted from samples using TRIZOL™ according to the manufacturer’s protocol. RNA Purity, concentration, and integrity were assessed using a NanoDrop One spectrophotometer (Thermo Scientific, Wilmington, DE, USA) and an Agilent Bioanalyzer 2100 system (Agilent Technologies, Santa Clara, CA, USA). All samples had a 260:280 nm ratio between 1.9 and 2.1 and RNA integrity number ≥ 9.5. Samples were sent to Novogene Corporation Inc. (Sacramento, CA, USA) for sequencing in Illumina platform. Data quality control was performed with FastQC v.0.11. After data filtering for adapters, clean reads were mapped to the C. elegans WS245 reference genome using HISAT 2.1.0 (81). Then, featureCounts v1.22.2 (82) was used to count the number of reads for each of the identified genes. Gene count matrix was filtered for genes with constant values (either 0 or empty). The gene counts were then loaded in RStudio for differential expression analysis with DESEQ2.0 (83). Counts were normalized using the negative binomial distribution under DESEQ2.0 and genes with log2 fold changes |> 1|, and false discovery rates FDRs < 0.05 were defined as Differential Expressed Genes (DEGs) and captured for further analysis. Principal components analysis (PCA), 3D PCA volcano plots and heatmaps were performed with R v3.6.1 packages in custom in house scripts. In order to identify potential transcription factors involved in the observed phenotypic changes in α3ΔN mutants, we utilized the JASPAR (84) database. JASPAR provides a comprehensive repository of DNA-binding profiles represented as position frequency matrices for transcription factors across multiple species. We specifically employed the JASPAR CORE database, which contains experimentally validated binding profiles for *C. elegans* (https://jaspar.elixir.no). Briefly we input our RNA-seq data and searched for transcription factors that show differential binding activity in the α3ΔN strain compared to the wild type. We analyzed the enriched transcription factor binding sites using the JASPAR enrichment tool, which facilitates the identification of significantly over-represented transcription factors in the genomic regions of interest. The results were visualized using custom scripts to plot the significance of transcription factor involvement, aiding in the mechanistic interpretation of the systemic changes observed in the omics data.

### Integration of proteomics with bulk RNA-seq through Projection to Latent Structures sparse-PLS

We utilized sparse Partial Least Squares (sPLS) in canonical mode via the mixOmics (47) R package to integrate high-dimensional proteomics and RNA-seq data. The plotIndiv function was used to visualize individual observations in a joint subspace, colored by condition groups, facilitating an assessment of data integration and clustering. Subsequently, the cim function was employed to create a Clustered Image Map (CIM), depicting the correlation patterns between both omic datasets. In addition, to identify key molecular drivers for each dataset, we analyzed the top contributors from the sPLS loadings. Results were visualized using plotLoadings, highlighting influential genes and proteins that significantly influence the observed correlation patterns latter translated into biological processes impacting both molecular layers confirmed subsequently with the Enrichment analysis.

### Enrichment Analysis

Gene Set Enrichment Analysis (GSEA) (41) was performed using the ClusterProfiler (85) library in-house implementation in R studio to assess enrichment signatures in the expression profiles in both TMT-MS and RNA-seq. The entire gene lists were pre-ranked based on the mean fold change and significance (FDR) of each gene. The analysis included the gene set lists from the curated Gene Ontology (GO) the proteostasis network consortium (48) and the chaperome subnetwork datasets (49). The significance of enrichment was set by Benjamini-Hochberg < 0.05. Regarding, chaperome subnetwork components enriched categories were identified by hypergeometric distribution FDR < 0.05.

### Toxicity assays

Toxicity assays were conducted following the guidelines of Qi Jia & Derek Sieburth (53) using stock solutions of juglone (50 mM in DMSO), tBoOH (70%, 7.2 M), and H2O2 (30%, 9.8 M), freshly prepared before each assay. Synchronized Day 1 adults (unless specified otherwise) C. elegans (a minimum of 40–100 worms) were placed in 1.5 mL Eppendorf tubes containing M9 buffer and washed three times. Oxidants were then added at final concentrations ranging from 50– 300 µM for juglone, and 100–2000 µM for both tBoOH and H2O2, followed by a 4-hour incubation on a rotating mixer. Post-incubation, worms were washed three times with M9 and transferred to fresh NGM plates seeded with OP50 for a 16-hour recovery period in the dark at 20°C. Survival rates were determined by counting alive versus dead worms. This procedure was repeated across various concentrations in triplicate for a minimum of two separate experiments. Mortality rates were analyzed using probit analysis through the BioRssay (86) package in R, which also estimated the lethal dose 50 (LD50), 95% confidence limits (CL), slope, and chi-square (χ2) values. Probit regression lines of log concentration against percent mortality facilitated the determination of LD50 values for each treatment, as detailed in Supplemental Figure 4B-E.

### Lifespan assays in xbp-1null hsp4::GFP worms

For lifespan experiments on xbp-1 null; hsp-4::GFP and α3ΔN; xbp-1 null; hsp-4::GFP strains, synchronized day-one adults were collected and seeded in OP50 NGM plates containing 100 µM FUdR (RPI) to inhibit reproduction. Animals were scored daily as dead if they exhibited no movement upon mechanical stimulation, and censored if death resulted from extraneous causes (e.g., drying out, internal hatching, or vulval protrusion). Worms were transferred to fresh plates every 2–3 days to prevent starvation. Statistical analyses were performed using the survminer and survival packages in R (https://github.com/kassambara/survminer/tree/master), and Kaplan–Meier curves were generated in GraphPad Prism version 8.0.0 (San Diego, CA, USA) (N ≥ 90). A p-value < 0.05 was considered statistically significant. All experiments were conducted at 20°C and repeated at least three times under blinded conditions.

### Toxicity assays Aldicarb

For Aldicarb toxicity assay a 10 mM stock solution of Aldicarb in DMSO was prepared, and synchronized day 1 adult worms were placed on NGM plates containing 1 mM Aldicarb (54). Survival rates were recorded every 15 minutes over the course of three independent experiments, with the data presented as means ± SEM. Statistical analysis of the assays was conducted using the survminer and survival packages in R as described above.

### Endoplasmic reticulum stress with tunicamycin

To induce ER stress we followed the guides of Gusarov et al. 2021 (87), C. elegans were placed on NGM plates supplemented with 50 µg/ml tunicamycin in a maximum of 2% DMSO, and compared to control groups on plates with only 2% DMSO. Treatment started on day 1 of adulthood, with worms being transferred to fresh plates every 3-4 days. Survival rates were monitored daily across three independent experiments, with results expressed as means ± SEM. Data analysis was conducted using the survminer and survival packages in R.

### Translation Assay, 6-FAM-dC-puromycin incorporation

To assess protein synthesis, we utilized a fluorescent puromycin incorporation method, employing 2 mM 6-FAM-dC-Puromycin (Jena Bioscience), according to Wang et al. 2013 (45) with some modifications for *C.elegans*. Day 1 adult worms were collected in M9 buffer, then rinsed and transferred into S-basal medium. An overnight OP50 bacterial culture was concentrated 10-fold in S-basal medium. Worms were subsequently placed into a mixture of 250 µL S-basal medium, 200 µL of the 10-fold concentrated OP50, and 5 µL of 2 mM 6-FAM-Dc-puromycin, also in S-basal. This mixture was incubated for 4 hours on a rotating mixer. Following incubation, worms were washed at least three times in S-basal, sonicated, and centrifuged for protein extraction. After sonication, two volumes of S-basal buffer were added, and the mixtures were centrifuged using Amicon Ultra Centrifugal Filters (YM-3, Millipore) to eliminate unincorporated 6-FAM-dC-Puromycin. A minimum of 10% of the reaction products were saved as input to quantify the total nascent protein chains using a plate reader. The incorporation rate was normalized against worms treated with 10 mM Cicloheximide (CHX) alongside 6-FAM-dC-Puromycin, to account for basal levels of protein synthesis inhibition.

### Degradation Assay

To evaluate the rates of time-dependent protein degradation, we utilized Cycloheximide (CHX). Day 1 adult worms were exposed to CHX (50 mg/ml in ethanol, Sigma-Aldrich) alone or in combination with the proteasome inhibitor MG132. We distributed and incubated groups of 800– 1000 worms each at 20°C for time intervals of 0, 1, 3, and 6 hours. Following treatment, we harvested the worms for total protein analysis. Protein was extracted and concentration was normalized through Bradford assay. The proteins were separated using NuPAGE 4-12% Bis-Tris gels (Invitrogen) and subjected to electrophoresis. Post-electrophoresis, gels were stained with Coomassie blue dye and then destained over a 24-hour period following standard protocols. To quantify degradation rates, we calculated the treated-to-untreated (CHX/CHX+MG132) protein ratio for each time point. This ratio was then normalized by division with the ratio at the initial time point (0 hours). Protein band intensities were quantified employing the Fiji Band/Peak Quantification Tool (NIH Image).

### Analysis of Heat stable Dirsorder proteins

Synchronized day 1 adult worm populations were collected in M9 Buffer and then diluted with 2× LDS Sample Buffer (Invitrogen). The samples were subjected to a controlled heat treatment to induce misfolding and precipitation of folded proteins following Park et al, 2022 (3), starting with an incubation in a 70°C water bath for 15 minutes, with vortexing every 5 minutes, followed by an increase to 90°C for 5 minutes. This thermal stress disrupts the normal folded states of proteins causing aggregation and precipitation of well folded proteins, leading to an increased presence of intrinsically disordered proteins, which lack folded structure and are this heat resistant. After the heat treatment, the samples were centrifuged at 6,000 rpm for 15 minutes at 4°C to remove any insoluble precipitates. The samples were normalized through Bradford assay and then loaded onto a NuPAGE 4-12% Bis-Tris gel (Invitrogen) for electrophoresis. Following electrophoresis, the gels were stained with Coomassie blue dye and destained over 24 hours according to standard protocols. Protein bands were quantified using the Fiji Band/Peak QuantificationTool (NIH Image).

### Confocal microscopy

#### Stress reporters hsp-4::GFP, hsp-6p::GFP and hsp-16.2p::GFP

Transcriptional reporter strains expressing hsp-4::GFP, hsp-6p::GFP, and hsp-16.2p::GFP, crossed into the α3ΔN background, were imaged using an EVOS M7000 microscope. Synchronized worms were cultured at 20°C until day 1 of adulthood, manually picked, and immobilized on slides with 25 mM sodium azide. Images were acquired within 5 minutes at 4× or 20× magnifications. For time-course experiments, synchronized worms were imaged at each developmental stage. Imaging was performed in three independent experiments with the following sample sizes: n = 438 for hsp-4::GFP, n = 428 for α3ΔN;hsp-4::GFP, n = 121 for hsp-6p::GFP, n = 123 for α3ΔN;hsp-6p::GFP, and n = 10 for each strain of hsp-16.2::GFP at day 1 adulthood. GFP fluorescence intensities were quantified using Fiji (NIH, version 2.1.4). A consistent threshold was applied to all images to create a selection, which was then quantified using the “Analyze → Measure” function in the ROI. In addition, images were captured under identical exposure settings to minimize technical variability. Data were plotted using Prism 8, and statistically significant differences were determined using two-way ANOVA or paired parametric t-tests.

#### vit-2::GFP and sGFP::ATZ

Defective mutants for accumulation of vitellogenin-2 vit-2::GFP outcrossed strain RT99 and the misfolded human protein, ATZ, involved in human α1-antitrypsin deficiency (ATD) strain were crossed into α3ΔNs. Synchronized worms were imaged throughout adulthood to assess GFP accumulation. For vit-2::GFP, images were captured using an EVOS M7000 microscope at 4× magnification, whereas for sGFP::ATZ, imaging was performed with a Zeiss LSM 710 multiphoton confocal microscope at 10× magnification. Worms were mounted on a 2% agar pad, anesthetized with 25 mM sodium azide, and covered with a glass coverslip for imaging. At least 21 worms per strain were quantified for sGFP::ATZ aggregates using a consistent threshold applied to all images to create a selection and then the “Analyze → Analyze Particles” function in Fiji was used. A minimum of 10 worms per strain were assessed per day for vit-2::GFP. The bright pharyngeal signal in sGFP::ATZ images is a genetic marker and was excluded from aggregate quantification. Images were processed and quantified in Fiji as described above, and data were plotted in Prism 8. Statistical differences between ATZ aggregates were determined using an unpaired t-test, whereas multiple t-tests (with the Benjamini, Krieger, and Yekutieli correction, FDR < 0.01) were applied to assess significant differences in vit-2::GFP levels. Each time point was analyzed independently without assuming uniform variance across tests.

#### Proteasome inhibition assay in vit-2::GFP

Dependency of proteasome degradation for vit-2::GFP was assessed by treatment with MG132 which completely inhibits WT and open-gate α3ΔN proteasomes. Worms were synchronized and similarly for toxicity assays were collected in S-basal and exposed to 50µm of MG132 for 1h, and 3h. Afterwards, worms were washed twice with S-basal, lysate and spun down for protein extraction. At least 20% of the total protein products were used as input to estimate the amount of vit-2::GFP signal in a plate reader. Experiments were performed at least three times independently and percentage of increase was obtained by diving the value of 1h and 3h by the starting value or time 0 of treatment. Significance was obtained with Unpaired t test with Welch’s correction.

#### Proteasome inhibition assay in sGFP::ATZ

The role of proteasome activity in the degradation of sGFP::ATZ aggregates was examined through MG132, an inhibitor for both WT and α3ΔN proteasomes. Worms were synchronized placed on FUDR plates and subsequently treated with MG132 at a concentration of 100 µM every 8 hours from day 2. DMSO-treated worms served as controls. On day 7, worms were collected and washed twice with M9 buffer. Then, worms were immobilized using 25 mM Sodium Azide on microscope slides and imaged using a Zeiss LSM 710 multiphoton confocal microscope at 10x magnification.

Image processing was performed as described above. To ensure reproducibility, these experiments were conducted at least three times independently. The number of aggregates was quantified in each condition and treatment. Statistical significance of the results was determined using one-way ANOVA, followed by Tukey’s post hoc test.

## Supporting information

DataS1-DEP

DataS2-DEG

DataS3-TFs-AnalysisResults

DataS4-Oxidation

DataS5-Strains & Alleles

## Acknowledgments

Our sincere appreciation extends to the Smith Lab members for their insightful discussions and feedback throughout the manuscript’s development. We thank Dr. Kevin Courtney and Dr. Leonardi for careful and meticulous review of our manuscript. We also acknowledge Dr. Neil Billington and the West Virginia University Microscope Imaging Facility for their support in conducting the confocal and imaging experiments.

## Author contributions

Conceptualization: DMS, RA, DST Methodology: DST, NA, RA, DMS Investigation: DST, NA, DMS Visualization: DST, NA Supervision: DMS, NA Writing—original draft: DST, DMS

Writing—review & editing: DST, DMS, NA

## Data and materials availability

All data are available in the main text or the supplementary materials. The sequencing data have been deposited at the National Center for Biotechnology Information’s Gene Expression Omnibus (GEO) and can be accessed via GEO Series accession number GSE259419. The mass spectrometry proteomics data have been deposited to the ProteomeXchange Consortium via the PRIDE partner repository with the dataset identifier PXD055640

Any additional data pertinent to this study are included within the article, supplementary files, or can be obtained from the corresponding author upon reasonable request.

## Supplementary Materials

Please see supplemental files Figure S1-S5 and Data S1-S5

