## Supplemental-Information for "Hyperactive 20S Proteasome Enhances Proteostasis and ERAD in *C. elegans* via degradation of Intrinsically Disordered Proteins"

David Salcedo-Tacuma *et al.*

**This PDF file includes:**

Figs. S1 to S5

**Other Supplementary Materials for this manuscript include the following:**

Data S1 to S5

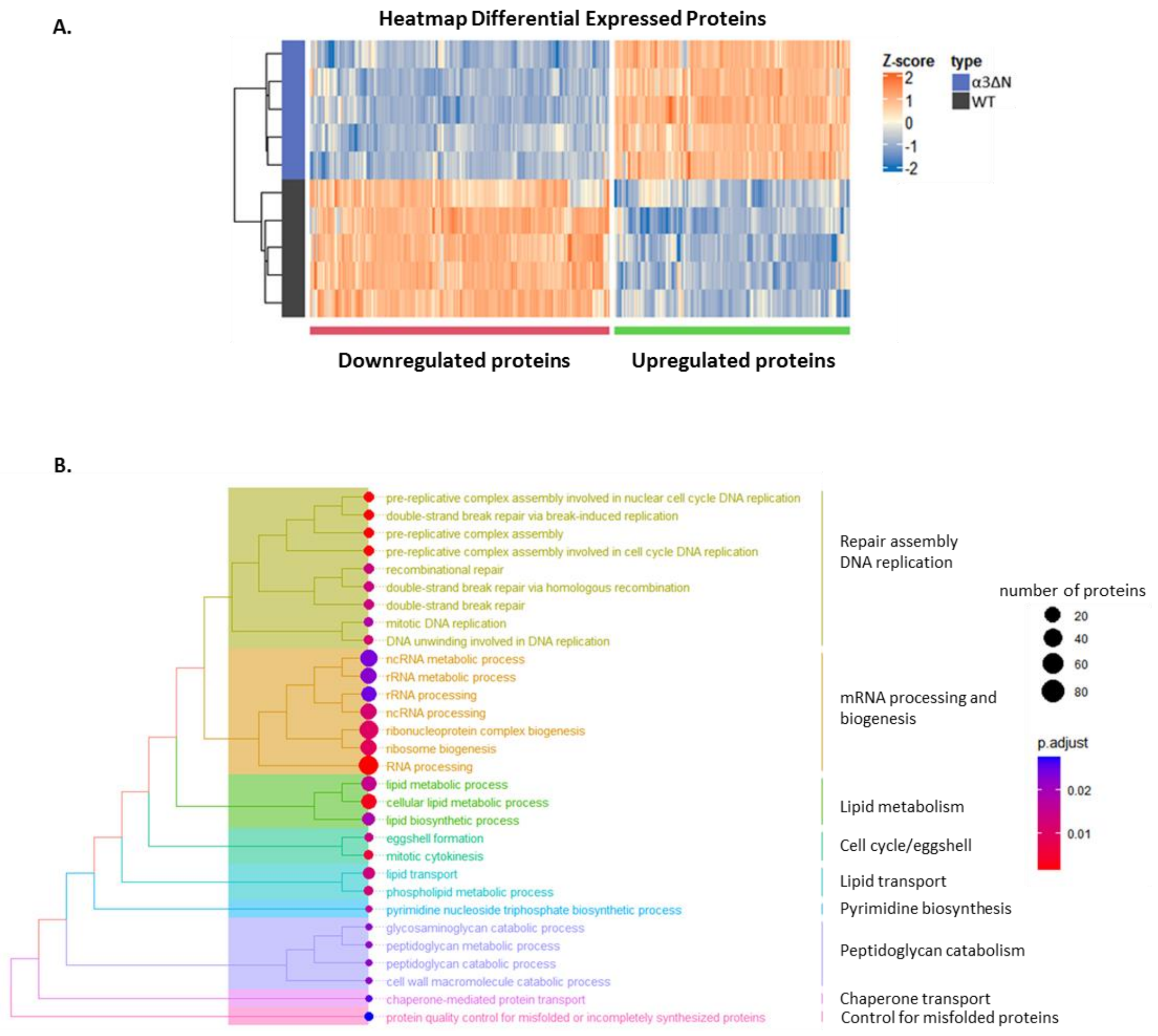


**Fig. S1.**

**Proteomic enrichment in α3ΔN mutants. A.** Heatmap of differentially expressed proteins (DEPs) in α3ΔN mutants. The hierarchical cluster above the heatmap distinguishes WT (black) from α3ΔN mutants (blue), using z-score values to indicate upregulation (orange) and downregulation (blue) (FDR<0.05) **B.** Treeplot of enriched biological functions from TMT-MS data, clustered by semantic similarity to illustrate the breadth of proteomic alterations in α3ΔN mutants. Seven color-coded clusters summarize key proteomic changes, with each branch ending in a dot representing the category's significance and size, as detailed in the adjacent panel; dot colors correlate with adjusted p-values for DEPs.

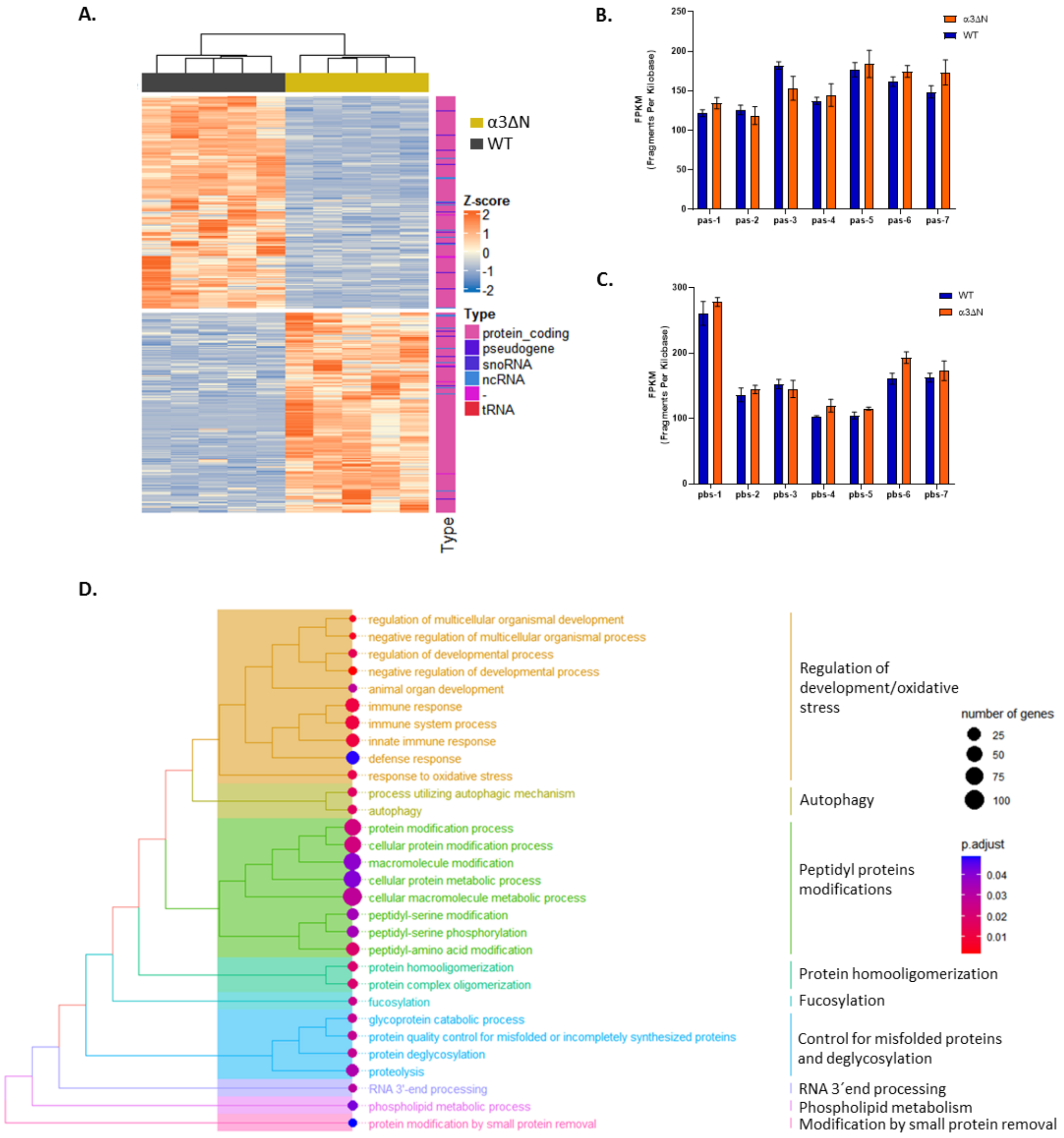


**Fig. S2.**

**Transcriptomic landscape and functional enrichment in α3ΔN mutants. A.** Heatmap visualization shows DEGs in α3ΔN mutants, with a side bar indicating transcript types through color coding, highlighting the predominance of protein-coding genes alongside ncRNAs and snoRNAs. The hierarchical clustering above the heatmap differentiates WT (black) from α3ΔN mutants (yellow), based on z-score values of expression: orange for upregulated and blue for downregulated transcripts (FDR<0.05). **B-C**. mRNA expression of proteasome subunit genes: Panel B presents FPKM-based expression levels of proteasome α-subunits, and Panel C details the expression of β-subunits in α3ΔN mutants, with data shown as mean ± SEM. No differential expression was observed among these genes. **D.** Treeplot of Transcriptomic Enrichment. Biological functions derived from RNA-seq data, organized by semantic similarity to reveal comprehensive transcriptomic alterations in α3ΔN mutants. Nine color-coded clusters summarize the principal changes observed, with each branch terminating in a dot that reflects the category's significance and size, as elaborated in the adjacent panel; dot colors correspond to adjusted p-values.

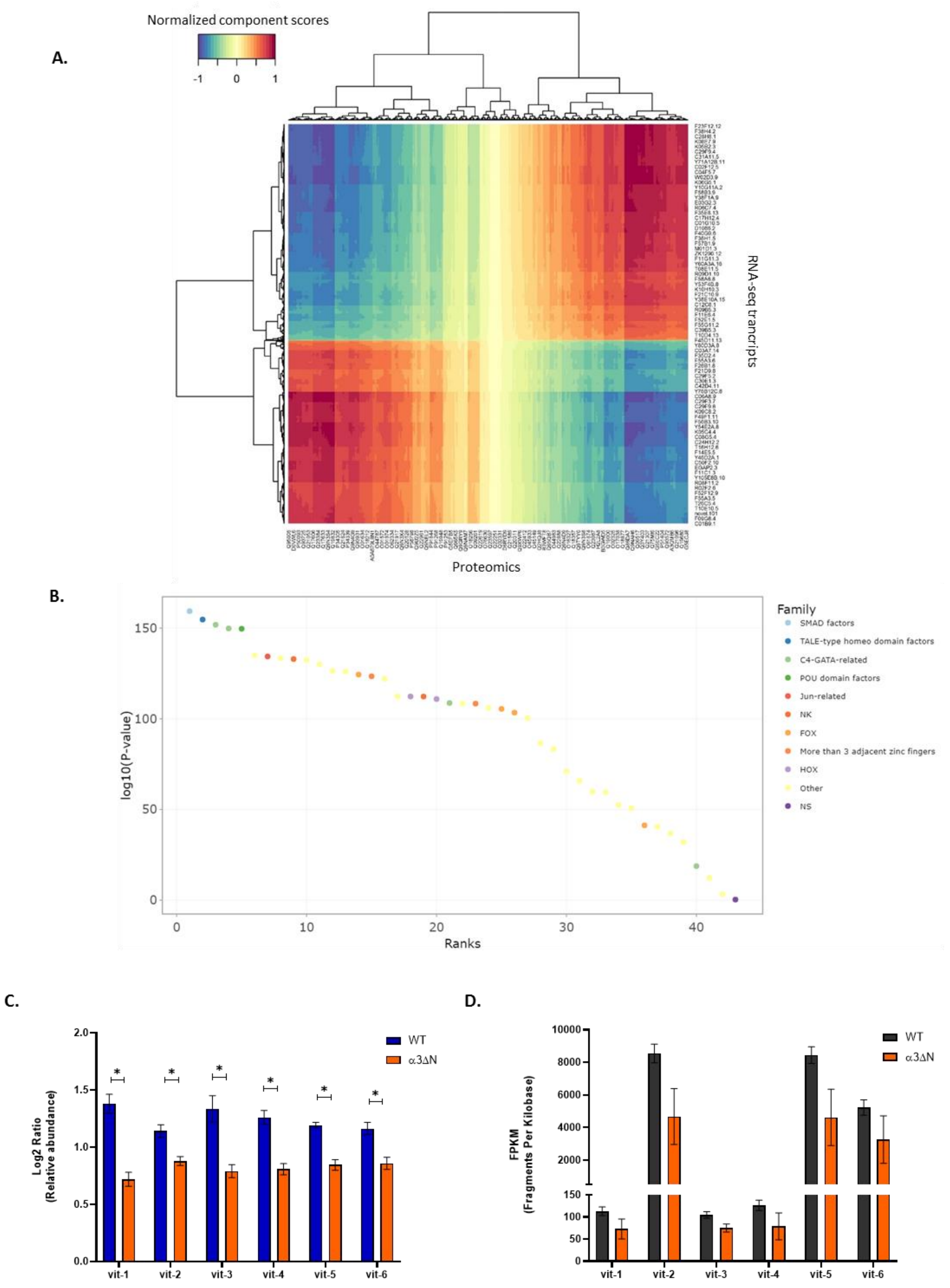


**Fig. S3.**

**Integration of proteomics and RNA-seq. A.** Clustered Image Map (CIM): This CIM visualizes the comprehensive correlation matrix between proteins and RNA transcripts, elucidating their interaction patterns. The clustering helps highlight groups of proteins and transcripts that co-vary significantly between the α3ΔN and WT conditions, suggesting potential regulatory mechanisms or pathways influenced by these conditions. **B.** Transcription Factor enrichment analysis with JASPAR. The X-axis represents the ranks assigned based on the statistical significance, with lower ranks indicating greater importance. The Y-axis shows the –log10 of the p-values. Each dot represents a transcription factor, color-coded by family to delineate different types of transcription factors involved, including SMAD factors, TALE-type homeo domain factors, and others (see TableS3). The analysis shows specific TF families that are enriched in the α3ΔN strain according to omics data, highlighting potential key regulators in the observed systemic adaptations. **. C-D.** Panel A details the log2 relative protein abundance of vitellogenins, indicating altered protein levels in α3ΔN mutants. Panel D presents vitellogenin gene expression measured in FPKM, highlighting no significant transcriptional changes. Both panels feature data as mean ± SEM, with asterisks (*) marking significantly different expressions (FDR < 0.05).

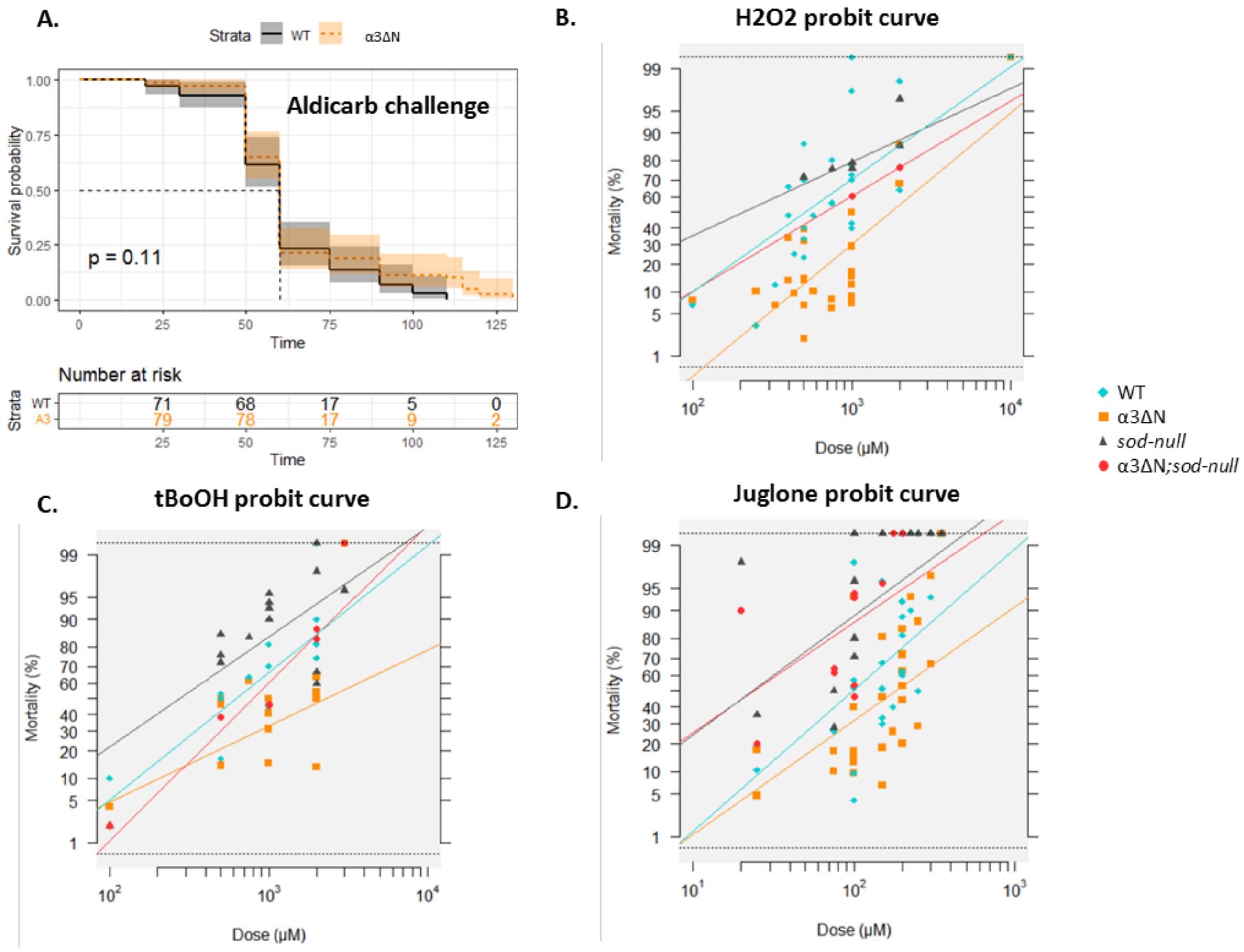


E.

| Compound | α3ΔN | | WT | | |
|---|---|---|---|---|---|
| | LD50 µM | RR50 | LD50 µM | RR50 | *p-value* |
| $H_2O_2$ | 1758 | 1.32 | 512 | 1 | 0.0025 |
| tBoOH | 2260 | 1.28 | 622 | 1 | 0.01667 |
| Juglone | 182 | 1,84 | 99 | 1 | 0.0125 |

| Compound | α3ΔN-/-SOD | | -/-SOD | | |
|---|---|---|---|---|---|
| | LD50 µM | RR50 | LD50 µM | RR50 | *p-value* |
| $H_2O_2$ | 674 | 1.32 | 207 | 0.4045 | 0.00179 |
| tBoOH | 797 | 1.28 | 277 | 0.4453 | 0.03492 |
| Juglone | 24 | 0.2429 | 23 | 0.236 | 0.96382 |

**Fig. S4.**

**Analyzing oxidative stress dose-response in α3ΔN mutants. A.** Survival Analysis under Aldicarb Stress. Compares survival rates of day 1 WT worms and α3ΔN mutants exposed to 1mM aldicarb, based on n = 3 independent experiments. Statistical significance assessed via log-rank test (No Significant). **B-D.** Probit analysis of Oxidative DoseStress-Response utilizing the BioRssays package (Day 1 adults). Probit transformation was used to linearize the dose–response curves, thereby enhancing the accuracy of LD50 estimations. These plots depict the relationship between probit-transformed mortality rates and log-dose exposure to H2O2, tBoOH, and Juglone. *Sod-null* mutants (gray triangles) serve as a susceptible reference; α3ΔN mutants (orange squares) exhibit significantly higher resistance compared to both WT (cyan diamonds) and the *sod-null* reference. Crosses between *sod-null* and α3ΔN (red dots) demonstrate a notable decrease in resistance. **E.** LD50 Determination and Resistance Ratios for α3ΔN mutants vs WT. Statistical significance was determined by ANOVA with Bonferroni-corrected binomial GLM tests, evaluates strain resistance differences. Clustered Image Map (CIM): This CIM visualizes the comprehensive correlation matrix between proteins and RNA transcripts, elucidating their interaction patterns. The clustering helps highlight groups of proteins and transcripts that co-vary significantly between the α3ΔN and WT conditions, suggesting potential regulatory mechanisms

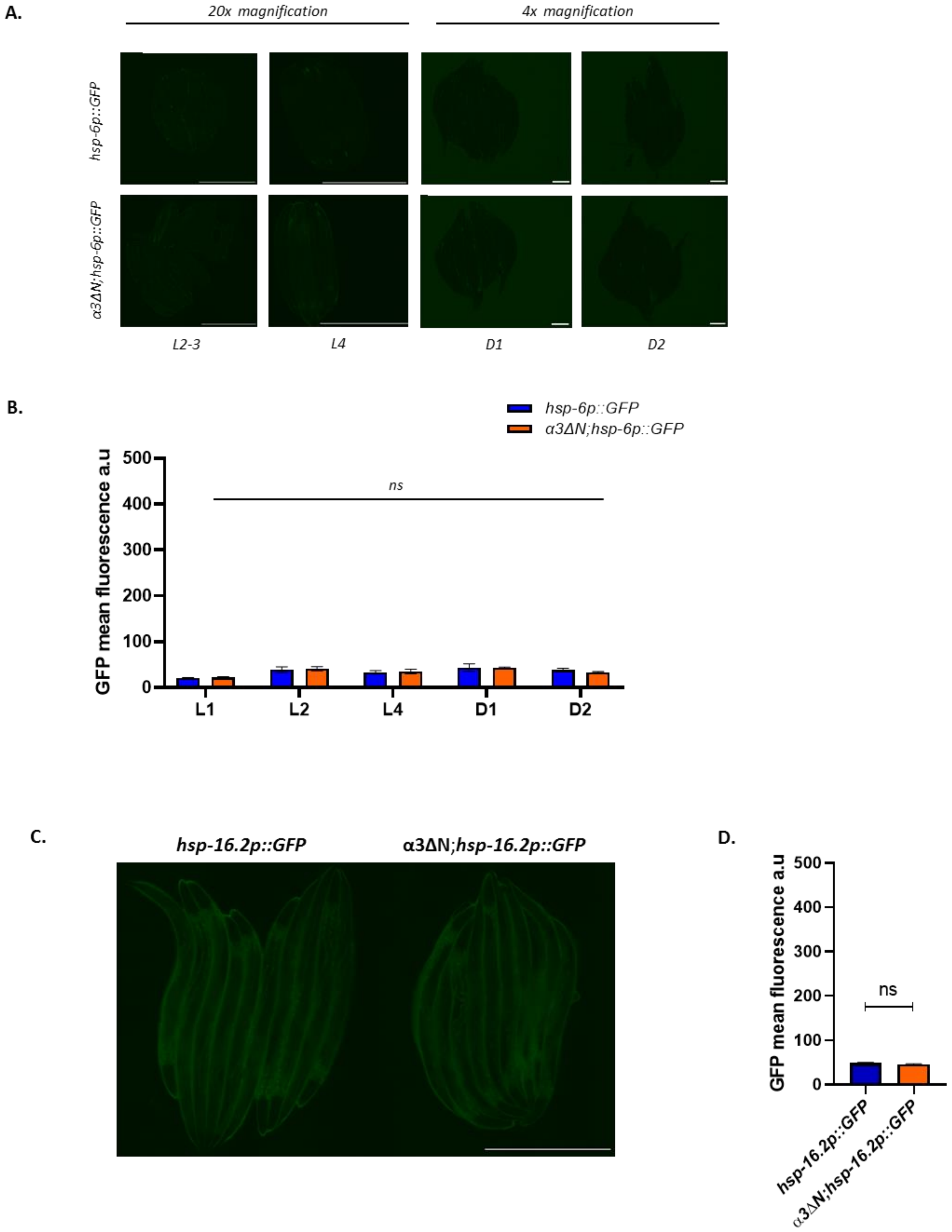


**Fig. S5.**

**Hyperactive proteasomes do not induce mitochondrial or cytosolic stress responses under basal conditions.**

**A.** Representative time-course images of the mtUPR reporter strain (hsp-6p::GFP, zcIs13) in both control and α3ΔN;hsp-6p::GFP worms under normal conditions. Early developmental stages were captured at 20× magnification, while adults were imaged at 4× magnification. **B.** Quantification of hsp-6p::GFP fluorescence intensity across early developmental stages and adulthood for both control and α3ΔN;hsp-6p::GFP worms. Data are presented as mean ± SD (scale bar: 500 µm). **C.** Representative images of the cytosolic heat shock reporter (hsp-16.2p::GFP, dvIs70) in control and α3ΔN;hsp-16.2p::GFP day 1 worms under basal conditions, captured at 20× magnification**. D.** Quantification of hsp-16.2p::GFP fluorescence intensity in day 1 adult worms, showing no significant induction of hsp-16.2 under normal conditions. Data are presented as mean ± SD (scale bar: 500 µm).

**Data S1. (separate file)**

DataS1-DEPs.xlsx. Differential Expressed Proteins (DEPs) detected.

**Data S2. (separate file)**

DataS2-DEGs.xlsx. Differential Expressed Genes (DEGs) detected.

**Data S3. (separate file)**

DataS3-TFs-AnalysisResults.xlsx. TF enrichment analysis results performed with JASPAR.

**Data S4. (separate file)**

DataS4-TFs-Oxidation.xlsx. Differential PTM Oxidation found in peptides detected by TMT-MS.

**Data S5. (separate file)**

DataS5-Strains and Alleles.xlsx. *C.elegans* Strains used in this study.
